# Serval: A modular framework for decoding imaging based spatial transcriptomics data

**DOI:** 10.1101/2025.10.09.681318

**Authors:** Jenkin Tsui, Naila Adam, Woongcheol Choi, Luna Y. Liu, Cristina Flores, Shadi Ansari, Esther Kong, Yukta Thapliyal, Hakwoo Lee, Shahid Haider, Isaac Von Riedemann, Ciara O’Flanagan, IMAXT Cancer Grand Challenges Consortium, Samuel Aparicio, Andrew Roth

## Abstract

Imaging-based spatial transcriptomics technologies have opened new avenues for studying cellular organization and gene expression within intact tissues. However, the accuracy of downstream analyses depends critically on the decoding step that reconstructs barcodes from fluorescence patterns and maps them to gene identities. Despite a growing number of decoding methods, systematic benchmarking has been limited. Here, we introduce Serval, a modular framework for developing and benchmarking decoding methods across diverse spatial transcriptomics platforms. Serval separates key decoding stages into independently configurable modules, enabling flexible integration of alternative algorithms. Using this framework, we develop CoSiD (Cosine Similarity Decoder), a novel method that improves transcript recovery by optimizing cosine similarity to known barcodes. We evaluate CoSiD and baseline methods on synthetic and real MERFISH datasets, showing that CoSiD achieves higher transcript recovery and superior correlation with expression references compared to existing methods. Furthermore, we demonstrate that the Serval framework generalizes beyond MER-FISH. By extending to the DART-FISH platform, we show that CoSiD improves transcript recovery, clustering stability and supports more direct annotation of complex biological structures such as the human primary motor cortex. These results establish that modular decoding frameworks facilitate robust, platform-agnostic benchmarking, ultimately supporting more accurate spatial transcriptomics analysis across diverse biological samples.

## Introduction

Imaging spatial transcriptomics (IST) maps gene expression while preserving tissue architecture, enabling quantitative analysis of how different cell populations are arranged and interact [1]. By knowing where genes are expressed in intact tissue, we can uncover principles of cellular organization, identify disease-associated domains, and reveal how microenvironments shape cell behavior. Imaging based spatial transcriptomics has already revealed spatial gradients of pathology in neurodegenerative disease [2], region-specific transcriptional programs in lung disorders [3], and tumor heterogeneity and resistant sub-populations when combined with single-cell RNA-seq [4]. Together, these studies highlight how spatial resolution in gene expression can drive major biological and clinical insights.

Despite the rapid advances of imaging based spatial transcriptomics methods, robust and accurate extraction of single-cell gene expression remains computationally challenging. A key computational step involves decoding multiplexed fluorescence barcodes, where accuracy directly determines downstream biological insights. Although numerous decoding methods exist, such as: MERlin [5], DeepCell-Spots [6], JSIT [7], and BarDensr [8], systematic benchmarking across different experimental scenarios is largely absent from current literature. This absence impedes methods selection and obscures comparative strengths and limitations.

Here, we present Serval, a modular framework that includes a re-implementation of MERlin and the Nearest Neighbor decoding strategies, providing a unified environment for benchmarking against external published pipelines. While recent efforts have focused on building standardized pipelines for FISH spatial transcriptomics [9], Serval addresses a complementary gap by introducing a framework for developing multi-step decoding pipelines which can include iterative fitting of parameters.

Using the Serval framework, we introduce CoSiD, which learns scaling factors by maximizing the cosine similarity between observed per-round fluorescence vectors and expected barcodes under regularization to suppress spurious signals. We evaluated CoSiD and baseline decoders on controlled synthetic datasets generated by a photorealistic simulator (varying signal-to-noise, transcript density, and background structure) in addition to 4T1 cell culture and implanted tumor MERFISH datasets. Across these tests, CoSiD yielded higher transcript recovery, improved correlation with external expression references, and reduced spurious detection of genes.

To demonstrate the cross-platform compatibility of Serval, we developed a new decoding workflow for the DART-FISH platform [10]. By reprocessing the human primary motor cortex DART-FISH dataset, we show that CoSiD improves transcript recovery and biological interpretability compared to the original sparse deconvolution approach (SpD). These results highlight Serval’s ability to facilitate robust decoding across distinct combinatorial imaging technologies.

## Results

### A Modular and Scalable Decoding Framework for Spatial Transcriptomics

Serval introduces a modular framework for developing decoding pipelines for multiplexed images. The design of Serval allows the decoding process to be separated into interchangeable components. To demonstrate the utility of the framework, we have implemented the MERlin [5] workflow for MER-FISH decoding. The MERlin workflow consists of several steps including: chromatic correction, image registration, image transformations, and spot detection (Figure 1).

**Fig. 1:**
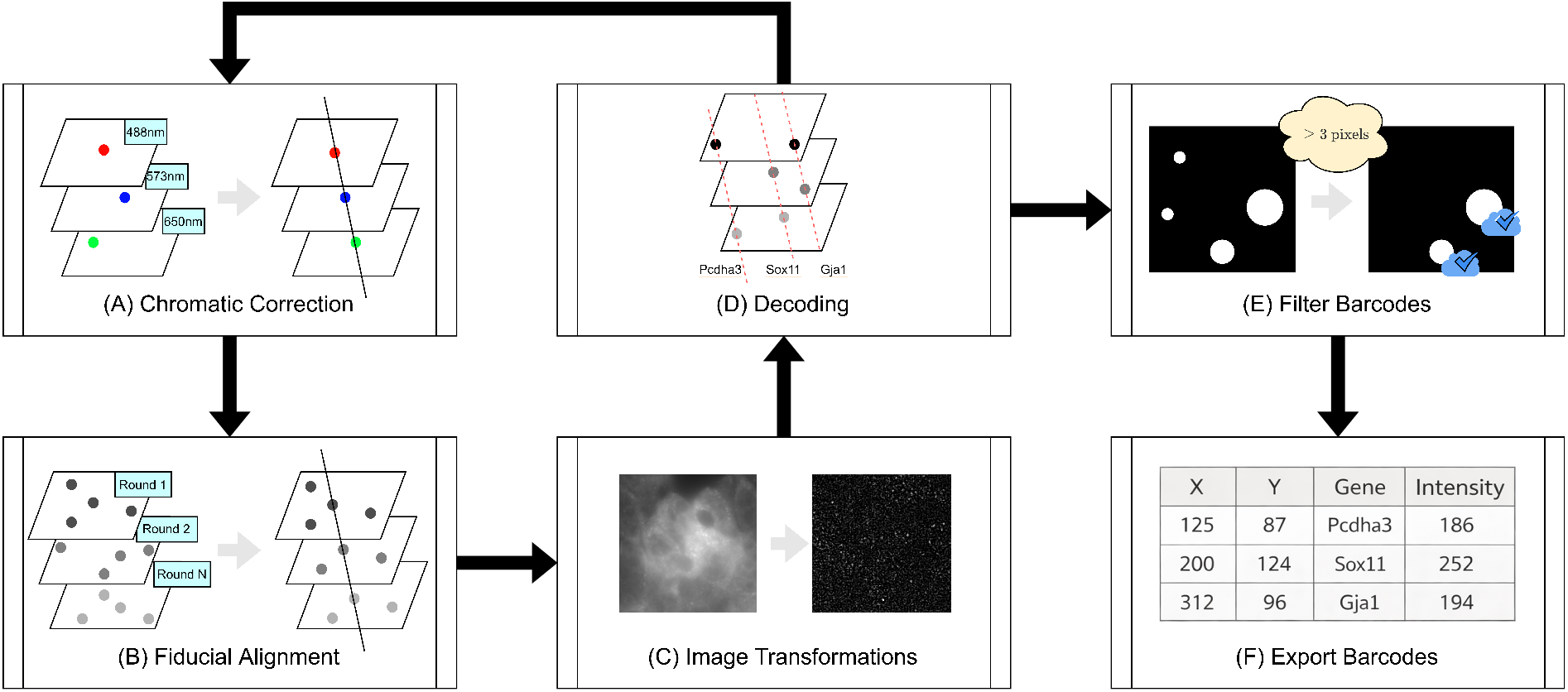
Serval MERlin decoding workflow. Serval’s MERlin pipeline comprises seven sequential modules: (A) chromatic correction to remove inter-channel misalignment; (B) fiducial alignment for sub-pixel registration across imaging rounds; (C) image transformations (background subtraction, denoising, deconvolution); (D) decoding of pixel intensities into barcode identities; (E) filtering of low-confidence or off-target barcodes; and (F) export of the final barcode table for downstream single-cell analysis.

The Serval implementation of the MERlin workflow separates decoding into distinct fit and predict stages. The fit stage estimates chromatic correction for each spot channel and scaling factors for each imaging round using a subset of the dataset. Serval leverages the Dask framework for out of core computation, allowing for memory efficient analysis of large datasets during the fit stage. The predict stage applies the learned parameters to decode barcodes and assign gene identities across the entire dataset. Each image stack, characterized by a field of view (FOV) number and a z-slice number, can be decoded independently using the fitted decoder and the corresponding scaling factors. This design allows decoding to scale efficiently across large datasets on compute clusters.

We applied both the Serval implementation and the standard MERlin pipeline to two real datasets: a cell culture sample and a tissue sample (Supplementary Figure 8). In the cell culture dataset, transcript counts per cell showed perfect agreement (Spearman = 1.000, *p*-value *<* 0.001) across 1017 cells (Supplementary Figure 8A). Similarly, in the tissue dataset, the two methods produced near-perfect concordance (Spearman = 0.999, *p*-value *<* 0.001) across 22705 cells (Supplementary Figure 8D). To further quantify absolute agreement, we computed Lin’s Concordance Correlation Coefficient [11]. For the cell culture dataset, Lin’s coefficient was 0.9999 (95% Confidence Interval: 0.9999–1.0000). For the tissue dataset, the coefficient was 0.9957 (95% Confidence Interval: 0.9956–0.9958).

We also performed a Bland–Altman analysis comparing transcript count per cell between the two methods. In the cell culture dataset, most of the points scatter within the 95% confidence interval band with no apparent trend, suggesting random noise (Supplementary Figure 8C). In the tissue dataset, we observed a positive bias and an upward trend at high transcript counts, indicating slight proportional bias and modest heteroscedasticity (Supplementary Figure 8F). Collectively, these results establish that the Serval implementation replicates the original MERlin implementation results to a high degree in both simple and complex sample contexts. In addition, the tissue results suggest that the Serval implementation may exhibit slightly greater sensitivity in heterogeneous tissue environments (Supplementary Figure 8B, Supplementary Figure 8E). We hypothesize that these small differences between Serval and original MERlin may arise from minor implementation-level numerical differences (e.g., optimization and floating-point operations), which become more apparent in heterogeneous tissue environments.

### A novel decoding algorithm maximizing cosine similarity

The original MERlin pipeline implements a heuristic decoding procedure that combines fiducial-based registration, deconvolution, and signal normalization. Barcode assignment is performed by minimizing the Euclidean distance between intensity vectors of candidate pixels and known barcode signatures. This distance is computed after vector normalization and per-round scaling. To learn the scaling factors, MERlin uses an iterative scheme where the initial scaling factors are used to decode the dataset. The detected spots from this step are then used to update the chromatic correction and round-specific scaling factors. This procedure is then iterated to update learned parameters, leading to an iterative conditional modes (ICM) like procedure. Specifically, we view the decoding of spots as an assignment of a latent variable to the highest-probability state (barcode or background). Conditioned on the decoded spots, the algorithm then updates the scaling factors. Surprisingly, the MERlin scaling factor update does not maximize the same objective used for decoding.

We reasoned that by aligning the objective between the two steps of the iterative procedure, we could improve the performance of the algorithm. To test this, we developed CoSiD, which finds scaling factors that maximize the cosine similarity between the decoded spots and the target barcode. To be precise, define the scaling vector **c** = [*c*_1_, *c*_2_, …, *c*_*d*_] (one scalar per imaging round), then CoSiD maximizes

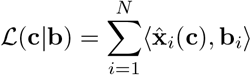

where 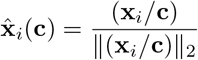 is the normalized scaled intensity vector for pixel *i* and **b** denotes the matched barcode vector obtained under the current scaling estimate. Following the ICM approach, the barcode assignments **b**_*i*_ are held fixed during the scaling optimization step.

In practice, direct optimization of the cosine objective can lead to pathological scaling factors resulting in over-representation of a single barcode. To address this we introduce a penalized objective function (see Supplementary Note 2: CoSiD Optimization for full gradient derivations).

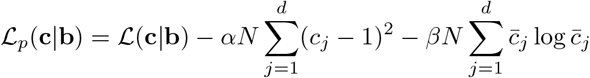

where *N* is the number of decoded (non-background) pixels used for optimization, *α* is the regularization coefficient for L2 penalty, *β* is the regularization coefficient for entropy and 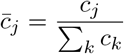 is the normalized scaling weight per round. The first term encourages pixel vectors to align with their matched barcodes. The L2 penalty anchors scaling values to 1, promoting signal consistency across rounds. The entropy penalty promotes diversity in scaling factors to prevent a few rounds from dominating. A detailed evaluation of parameter sensitivity and practical guidance for selecting these regularization weights across different datasets is provided in Supplementary Note 3: Practical Guidance for Selecting Regularization Parameters.

The proposed penalization scheme attempts to enforce uniformity across scaling factors, and avoids large values. In cases where there is significant variation in signal intensity between imaging rounds, for example due to different colour channels, this scheme can perform poorly. To address this issue we implement a pre-scaling step for the pixels. Specifically, we estimate a scaling factor **c**_0_ before model fitting. We substitute the raw pixel values **x**_*i*_ with their scaled values 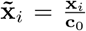. To estimate **c**_0_ we use the same strategy as MERlin uses to initialise scaling factors, namely setting **c**_0_ to the 90th percentile intensity for each imaging round across all FOVs.

We implemented CoSiD in Serval and created a new decoding pipeline by replacing the decoding step of the MERlin pipeline. All other steps of the pipeline were kept the same to allow us to isolate the impact of altering the decoding strategy.

### CoSiD accurately recovers transcript identity and abundance across diverse simulated imaging conditions

To assess decoder performance, synthetic images were generated using our multiplexed spot simulator across ten benchmarking scenarios (100 replicates per scenario). Collectively, these scenarios span the principal factors that influence transcript decoding accuracy in multiplexed imaging experiments, including signal strength (photon counts), signal-to-cell-background ratio (SCR), barcode corruption through bit drop and bit add events, nonspecific background fluorescence, round-to-round intensity variation, and complete round failure. By systematically varying these parameters individually and in combination, the benchmark captures a diverse range of realistic and adverse imaging conditions encountered in practice. Details of each scenario are provided in Supplementary Table 1.

Decoder performance was evaluated using two complementary classes of metrics: transcript-level detection accuracy and abundance recovery. Transcript-level detection accuracy was assessed using exact recall, exact false discovery rate (FDR), and exact F1 score, which quantify the recovery, precision, and overall accuracy of individual transcript detections relative to ground truth. Abundance recovery was assessed using the maximum Pearson correlation coefficient between decoded and ground-truth transcript abundances across threshold values. See Supplementary Materials (Decoder Benchmarking and Metric Computation on Synthetic Datasets) for a comprehensive description of the benchmarking framework and metric optimization pipeline.

The simulation results presented in Figure 2 focus on exact recall, exact FDR, exact F1 score, and maximum Pearson correlation coefficient. Additional performance metrics, including maximum Spearman correlation coefficient, exact average precision, localization recall, and localization F1 score, are provided in the Supplementary Materials (Supplementary Figure 1) to provide a more comprehensive evaluation of decoder performance.

**Fig. 2:**
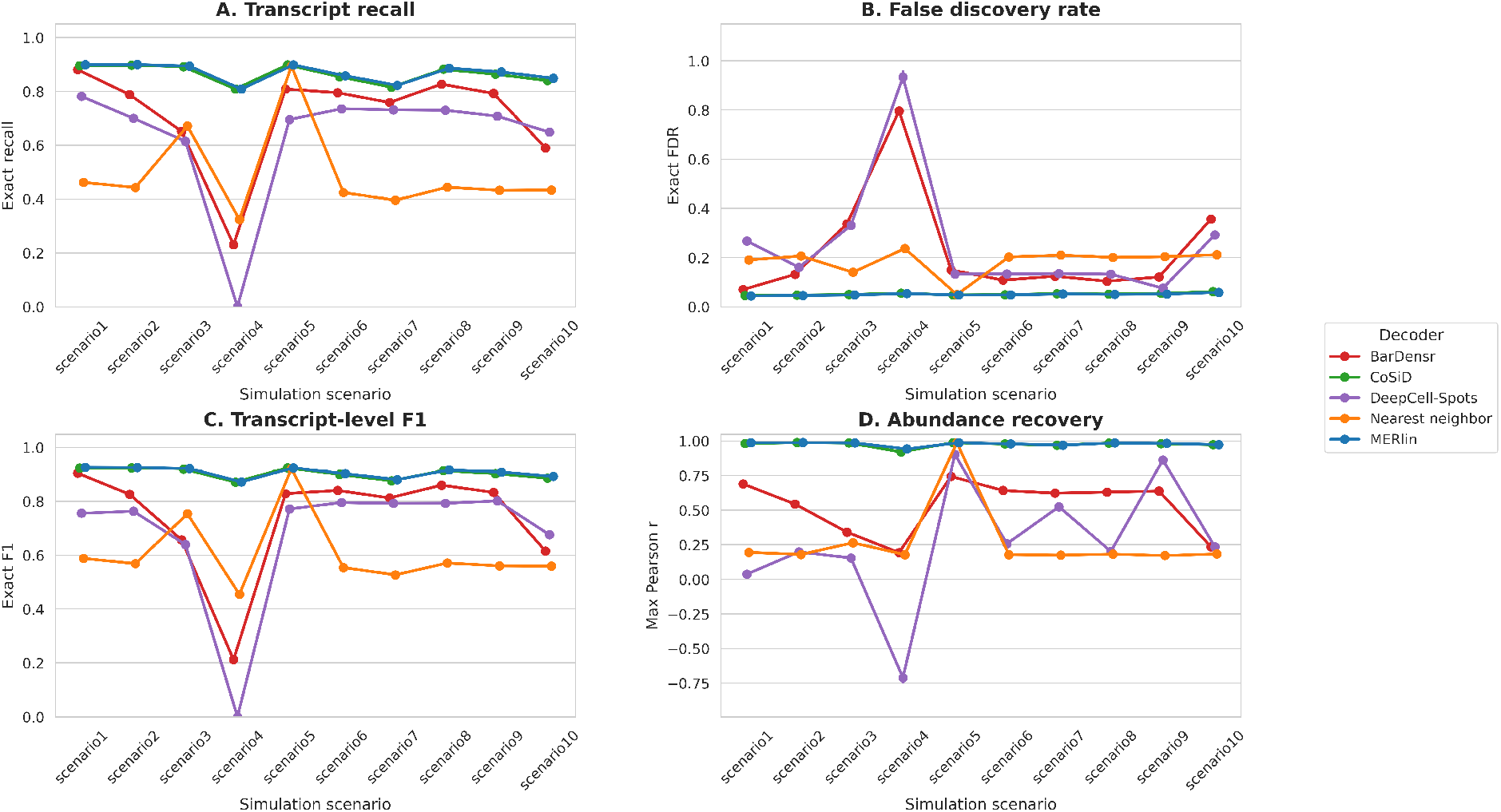
CoSiD matches MERlin performance and exceeds competing decoding methods in synthetic benchmarks. **(A)** Exact recall across the ten simulation scenarios defined in Supplementary Table 1. **(B)** Exact false discovery rate (FDR). **(C)** Exact F1 score. **(D)** Maximum Pearson correlation coefficient between decoded and ground-truth transcript abundances. Points represent the mean across 100 replicates per scenario and error bars indicate the standard error of the mean. Lines connect each decoder across scenarios to aid visual tracking of performance and do not imply continuity or ordering between scenarios. Decoder performance is shown for CoSiD (orange), MERlin (purple), BarDensr (blue), Nearest Neighbor (red), and DeepCell-Spots (green).

We compared CoSiD to the Serval implementation of MERlin algorithm and a baseline Nearest Neighbors algorithm which omits the scaling factor correction of CoSiD and MERlin decoders. We also compared CoSiD to the decoders in existing literatures, including: BarDensr and DeepCell-Spots. While JSIT was evaluated on the experimental datasets, it was excluded from the synthetic benchmark. JSIT requires extensive manual hyperparameter tuning and visual inspection for individual fields of view, rendering it computationally intractable for a high-throughput, automated simulation framework consisting of 1,000 independent replicates. CoSiD was run with L2 penalty of 0.001 and entropy penalty of 0.01.

To assess decoder performance while satisfying the assumption of independent measurements, statistical evaluations were conducted independently for each of the ten simulation scenarios. Within each scenario, decoder performance was compared using a Friedman omnibus test, utilizing the 100 stochastic replicates as independent blocks (*n* = 100). To control the family-wise error rate across the multi-metric evaluation within a given scenario, Friedman *p*-values were adjusted using a Bonferroni correction for the six evaluated metrics (significance threshold: adjusted *p <* 0.01). For scenario-metric combinations where the Friedman test indicated significant variance, post-hoc pairwise comparisons were performed using the Nemenyi test, with *p*-values further adjusted across all pairwise decoder combinations [12].

For the transcript-level detection metrics, CoSiD achieved a mean exact recall of 0.864, mean false discovery rate (FDR) of 0.051, and mean exact F1 score of 0.904 across the ten simulation scenarios (Figure 1A–C). MERlin achieved comparable mean recall (0.868), FDR (0.050), and F1 score (0.907). BarDensr, Nearest Neighbor, and DeepCell-Spots exhibited lower transcript-level performance, with mean recall values of 0.712, 0.492, and 0.634, respectively, and mean F1 scores of 0.739, 0.606, and 0.679, respectively (Supplementary Table 2).

For abundance recovery, CoSiD achieved a mean maximum Pearson correlation coefficient of 0.975, comparable to MERlin (0.978) and substantially higher than BarDensr (0.528), Nearest Neighbor (0.270), and DeepCell-Spots (0.267) (Figure 1D; Supplementary Table 2). Similarly, CoSiD achieved a mean maximum Spearman correlation coefficient of 0.972, comparable to MERlin (0.975) and higher than BarDensr (0.558), Nearest Neighbor (0.344), and DeepCell-Spots (0.722) (Supplementary Table 2).

The performance of decoders was significantly different across all ten scenarios and all six evaluated metrics (60/60 tests, adjusted *p <* 0.01, Friedman test). CoSiD and MERlin decoders did not show statistically different performance in approximately 80% of all pairwise evaluations (47/60 tests, adjusted *p >* 0.01, Nemenyi post-hoc test). Both decoders significantly outperformed the other methods: BarDensr, Nearest Neighbor, and DeepCell-Spots, in over 98% of all evaluated scenario-metric conditions (356/360 tests, adjusted *p <* 0.01, Nemenyi post-hoc test).

Overall, CoSiD and MERlin exhibited similar performance across the simulated benchmark, with both methods consistently outperforming BarDensr, Nearest Neighbor, and DeepCell-Spots across a broad range of imaging and decoding conditions. However, decoding performance varied across simulation scenarios, reflecting the distinct imaging challenges represented by each benchmark. In particular, the performance gap between CoSiD/MERlin and the alternative methods became more pronounced under lower signal-to-cell-background ratio (SCR), suggesting that accurate estimation of per-round scaling factors becomes increasingly important for distinguishing true barcode signals from background fluorescence.

### CoSiD achieves a superior correlation–yield tradeoff in raw spot decoding on the 4T1 MERFISH dataset

Having established that CoSiD matches MERlin on synthetic benchmarks, we next evaluated decoding performance on experimental data using the 4T1 cell culture and tissue MERFISH datasets. To isolate core decoding accuracy prior to cellular segmentation or background-gene filtering, we examined the correlation between decoded and reference transcript abundances (see Supplementary Methods for Global Expression Concordance) as a function of transcript yield, sweeping each framework across its stringency thresholds (see Supplementary Methods for Performance Evaluation on MERFISH Experimental Datasets).

Decoding accuracy and transcript yield trade off against one another: stringent thresholds recover few, high-confidence transcripts, whereas permissive thresholds recover more transcripts while admitting progressively more noise. Summarizing a decoder by the single highest correlation along this curve is therefore misleading, because the maximum correlation frequently falls at a low-yield operating point that recovers too few transcripts for reliable downstream analysis. The resulting sparse cell-by-gene matrices produce unstable clustering and cell type annotation. The operationally relevant criterion is instead the correlation a decoder sustains at the high transcript yields needed for downstream analysis, i.e., the right-hand end of the correlation–yield curve.

Judged on this basis, CoSiD achieved the optimal combination of correlation and yield in both datasets. In the cell culture dataset, CoSiD maintained a higher reference correlation than MERlin and Nearest Neighbor across the sweep. Most importantly, CoSiD sustained near-peak correlation out to the highest transcript yields (Figure 3A, top). Nearest Neighbor, by contrast, achieved competitive correlation only at low yield and did not reach comparable detection counts, demonstrating how a high-correlation focus can return a low-yield setup that offers limited downstream value. In tissue, CoSiD and MERlin were closely matched in correlation, but CoSiD extended to substantially higher transcript yields at equivalent correlation, recovering more usable transcripts without sacrificing correlation (Figure 3A, bottom). Across both datasets, CoSiD substantially outperformed the probabilistic and machine-learning frameworks BarDensr, DeepCell-Spots, and JSIT, which rely on native spot-confidence or intensity scores rather than distance thresholds (Supplementary Note 4: Parameter Sweep for Benchmark Baselines). Together these results indicate that CoSiD isolates true signal from raw pixel intensities at least as effectively as MERlin, and more effectively than competing methods, precisely in the high-recovery regime on which downstream analysis depends.

**Fig. 3:**
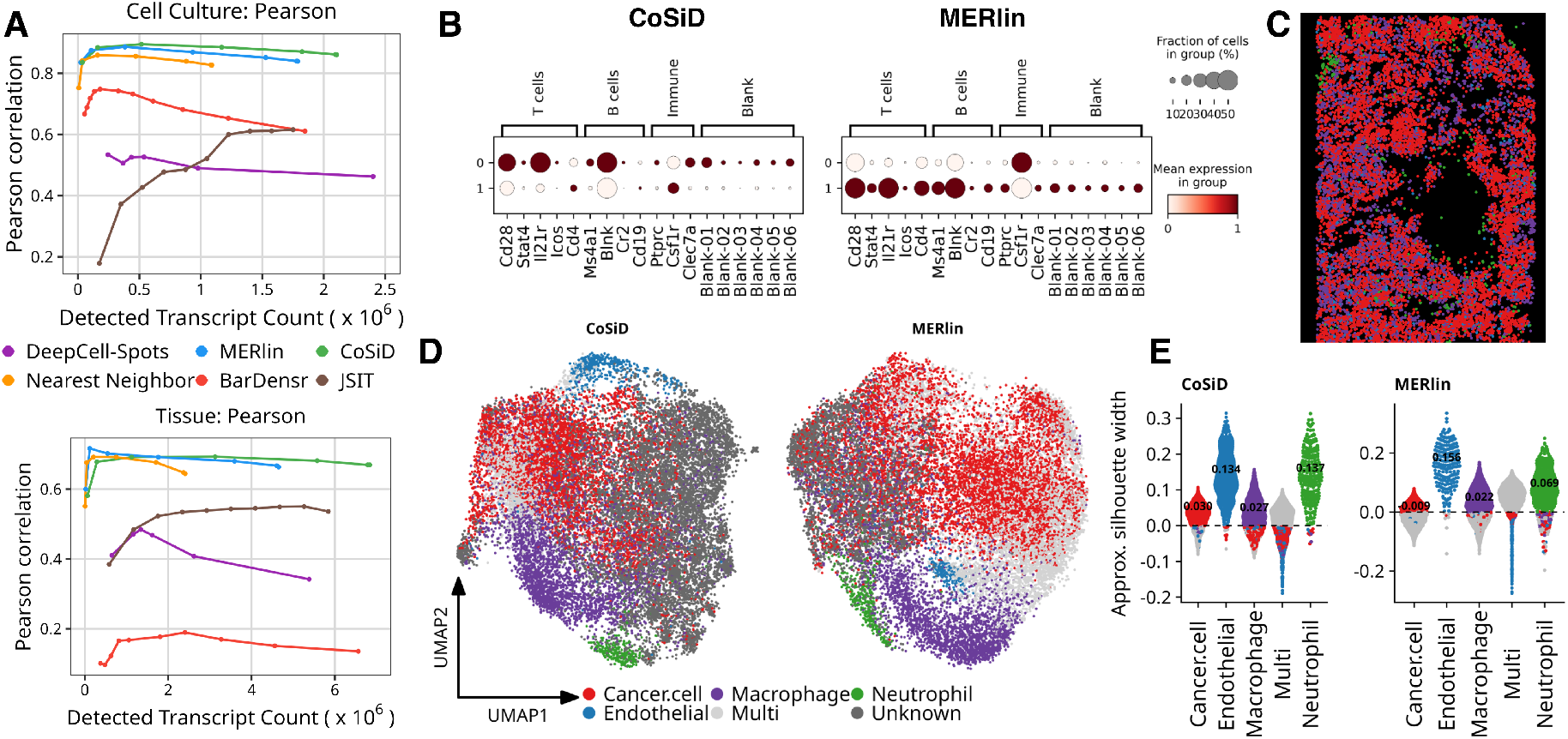
CoSiD improves transcript quantification and cell-type annotation coherence in the 4T1 MERFISH dataset. **(A)** Pearson correlations as a function of detected transcript counts in the 4T1 cell culture dataset (upper) and 4T1 tissue (lower), comparing six decoding methods (MERlin, Nearest Neighbor, CoSiD, BarDensr, DeepCell-Spots and JSIT). **(B)** Dot plots of off-target immune marker genes and blanks for CoSiD-decoded (left) versus MERlin-decoded (right) clusters from the cell culture dataset. **(C)** Spatial distribution of cell type annotations shown in (D). **(D)** UMAP visualization of CoSiD vs MERlin decoded 4T1 tissue, coloured by scGate annotation. ‘Unknown’ (dark grey) denotes cells lacking sufficient marker signal to pass any gate; ‘Multi’ (light grey) denotes cells with detectable but conflicting lineage signatures. **(E)** Approximate silhouette width by cell type for CoSiD and MERlin decoders. Median silhouette width per cell type is overlaid.

Mapping decoded transcripts onto segmented single-cell boundaries reproduced this advantage over the MERlin baseline. CoSiD recovered significantly more total transcripts per cell in both the cell culture (Cohen’s *d* = 1.51, *n* = 1,017 cells; Supplementary Figure 6A) and tissue (Cohen’s *d* = 1.04, *n* = 22,705 cells; Supplementary Figure 7A) datasets, with correspondingly higher transcript density per unit cell area (*d* = 2.04 and *d* = 1.29, respectively; all *p <* 0.001, Wilcoxon signed-rank test). This gain was accompanied by a reduction in the number of unique genes detected per cell (*d* = −0.63 in cell culture; Supplementary Figure 6B, *d* = −1.66 in tissue; Supplementary Figure 7B). This improvement in sensitivity was uniform rather than region-driven: CoSiD yielded higher transcript counts than MERlin in every field of view, in both the cell culture (15/15 FOVs, *d* = 2.44; Supplementary Figure 6D) and tissue (54/54 FOVs, *d* = 2.30; Supplementary Figure 7D) samples.

### CoSiD reduces spatial noise and improves downstream cellular annotation of 4T1 MERFISH dataset

A key motivation for increased transcript recovery is improving the cell-by-gene matrix used for down-stream transcriptomic analysis. Targeted spatial transcriptomics platforms measure restricted panels and involve more upstream steps, which amplify noise, making annotation more sensitive to missing transcripts and off-target signal. As a result, noise in the decoded expression matrix obscures biologically meaningful transcriptional structure, limiting the reliability of both reference-based integration with scRNA-seq and *de novo* cell-type identification. We therefore evaluated whether CoSiD improves the annotation-ready expression matrix against the MERlin baseline using the 4T1 MERFISH dataset (Figure 3, Supplementary Figure 10).

Our initial experiments included a 4T1 cell culture dataset, where CoSiD modestly reduced off-target T/B-cell marker gene signal relative to MERlin, with fewer cells expressing at least one T/B-cell marker gene(65.5% versus 74.0%; Figure 3B; Supplementary Table 7). However, the limited biological heterogeneity of the cell culture constrained further downstream analyses (Supplementary Figure 10B– D). Analysis of a reference 4T1 scRNA-seq dataset showed that cells were primarily separated by cell-cycle phase, and only a small number of discriminatory cell-cycle genes were represented in the targeted MERFISH panel. We therefore used the cell culture dataset as a technical specificity benchmark and focused downstream annotation analyses on the more heterogeneous 4T1 tissue dataset.

CoSiD produced a gene expression matrix with reduced off-target signal and improved overall spatial specificity metrics (Supplementary Table 3, Supplementary Table 4, Supplementary Table 5, Supplementary Table 6, Supplementary Table 7). Although CoSiD recovered more gene molecules than MERlin (5,878,085 versus 4,139,278), it produced far fewer blank barcode detections (1,020 versus 4,801), resulting in an estimated global FDR of 3.87 × 10^*−*5^ compared with 2.59 × 10^*−*4^ for MERlin (Supplementary Table 6). This represents an approximately 6.7-fold reduction in global decoding noise. Consistent with this reduction in off-target signal, the CoSiD count matrix exhibited higher sparsity (0.819 versus 0.723) and lower Shannon entropy (1.170 versus 1.665) than MERlin (Supplementary Table 3), indicating that recovered counts were concentrated on a smaller set of biologically relevant transcripts rather than distributed across off-target genes and blanks.Notably, this increase in specificity was achieved despite a higher transcript yield per cell (280 versus 198), reflecting improved signal quality rather than reduced recovery. Orthogonal control-gene analyses showed a similar pattern. In severely immunocompromised NRG tissue, where T- and B-cell markers are expected to represent off-target signal, CoSiD reduced the off-target marker fraction from 0.0064 to 0.0009 and the fraction of cells expressing at least one of these markers from 57.47% to 18.4% (Supplementary Table 5). CoSiD also increased signal-to-noise ratios (SNR) relative to both blank barcodes and T/B-cell marker controls (Supplementary Table 5, Supplementary Table 7). Additionally, CoSiD reduced biologically implausible co-expression as reflected by the mutually exclusive marker co-expression rate (MECR) metric, especially for marker pairs expected to be more distinct (e.g., Neutrophil–Macrophage) (Supplementary Table 5). Apparent co-expression can arise from segmentation ambiguity, transcript assignment errors, or spatially adjacent cell types; therefore, MECR is a limited specificity check constrained by marker panel and can only be interpreted in relative context.

To reduce dependence on clustering, which can vary with the underlying low-dimensional embedding, we annotated major cell classes using a marker-based scGate multiclassifier [13] framework rather than assigning labels directly from unsupervised clusters (Supplementary Figure 10A, Supplementary Figure 11A, B and F). Notably, under MERlin, marker signal is less restricted to target cell type with more cells co-expressing markers of multiple lineages at similar levels (Supplementary Figure 11F) which underlies the classification differences observed (Supplementary Figure 11A). We then quantified the coherence of these annotated cell classes in the expression-derived PCA space using silhouette width (Supplementary Figure 11C-E). Overall, CoSiD produced consistent improvements in annotation structure for several major populations (Figure 3C-E and Supplementary Table 4). The largest gains were observed for cancer cells and neutrophils; cancer cells shifted from a negative median silhouette width under MERlin to a positive value after CoSiD, with the fraction of cells showing negative silhouette widths decreasing from 59.2% to 20.2%. Neutrophils also showed improved separation, with median silhouette width increasing from 0.069 to 0.137 and the fraction of negative silhouette widths decreasing from 20.9% to 3.9% (Supplementary Table 4). Macrophage and endothelial annotations were broadly comparable between decoders, suggesting that the benefit of CoSiD was most evident for populations whose annotation was more sensitive to off-target signal or sparse marker recovery. Together, these results indicate that CoSiD improves the annotation-readiness of the 4T1 MERFISH cell-by-gene matrix, although the magnitude of improvement is modest in this dataset and likely constrained by panel design and tissue structure.

### CoSiD generalizes to DART-FISH technology enhancing clustering and integrative analyses

To test whether CoSiD improves downstream analysis in a distinct combinatorial imaging platform, we reprocessed the human primary motor cortex DART-FISH dataset. DART-FISH differs from MERFISH in both chemistry and image formation, using padlock-probe capture, rolling-circle amplification, and pixel-level sparse deconvolution (SpD) to decode amplified rolonies. In addition, a well-annotated snRNA-seq reference is available for cell-type annotation and label transfer [14].

For downstream DART-FISH analyses, we restricted the cell-by-gene matrix to decoded transcripts located within nuclear segmentation masks. Restricting transcripts to nuclear masks provides a conservative matrix that more closely matches the snRNA-seq reference, reducing the co-detection rate. Moreover, transcript-to-cell assignment in the released DART-FISH processing workflow includes a permissive maximum rolony-to-cell distance parameter (150 pixels), which upon examination proved to be very noisy and uninterpretable (Supplementary Figure 12). Overall, CoSiD performed comparably to MERlin on this dataset (41.4 vs 41.5 transcripts per nucleus; entropy 0.960 vs 0.964; sparsity 0.836 vs 0.833), with both recovering substantially more signal than SpD, as summarized in Supplementary Table 8 (25.5 transcripts per nucleus, sparsity 0.921, entropy 0.571). Interestingly, this contrasts with the MERFISH tumour data, where CoSiD diverged more clearly from MERlin. Since CoSiD differs from MERlin mainly in scaling-factor estimation, which matters more under lower signal-to-cell-background conditions as reflected in our simulations, we hypothesize that the two converge in cleaner, well-structured tissue such as this normal brain dataset and differ more in denser, more heterogeneous tumour tissue.

To avoid relying solely on unsupervised clusters, we generated major class annotations using two independent approaches: Seurat label transfer from the matched snRNA-seq reference [14] and marker-based scGate classification. High-confidence cells were then defined as cells receiving concordant major-class assignments from both approaches. CoSiD produced a well-resolved low-dimensional embedding with coherent, contiguous cluster topology, whereas SpD yielded a fragmented manifold with unstable, over-partitioned clusters difficult to annotate at equivalent resolution (Figure 4A; Supplementary Figure 12). Moreover, approximate silhouette width computed across major cell classes on high-confidence cells showed higher median silhouette widths for both excitatory (0.047) and inhibitory (0.013) neurons for CoSiD relative to SpD (0.008 and −0.058, respectively) (Supplementary Figure 13F). Thus, increased transcript recovery translated into a more interpretable low-dimensional structure and a more accessible annotation workflow.

**Fig. 4:**
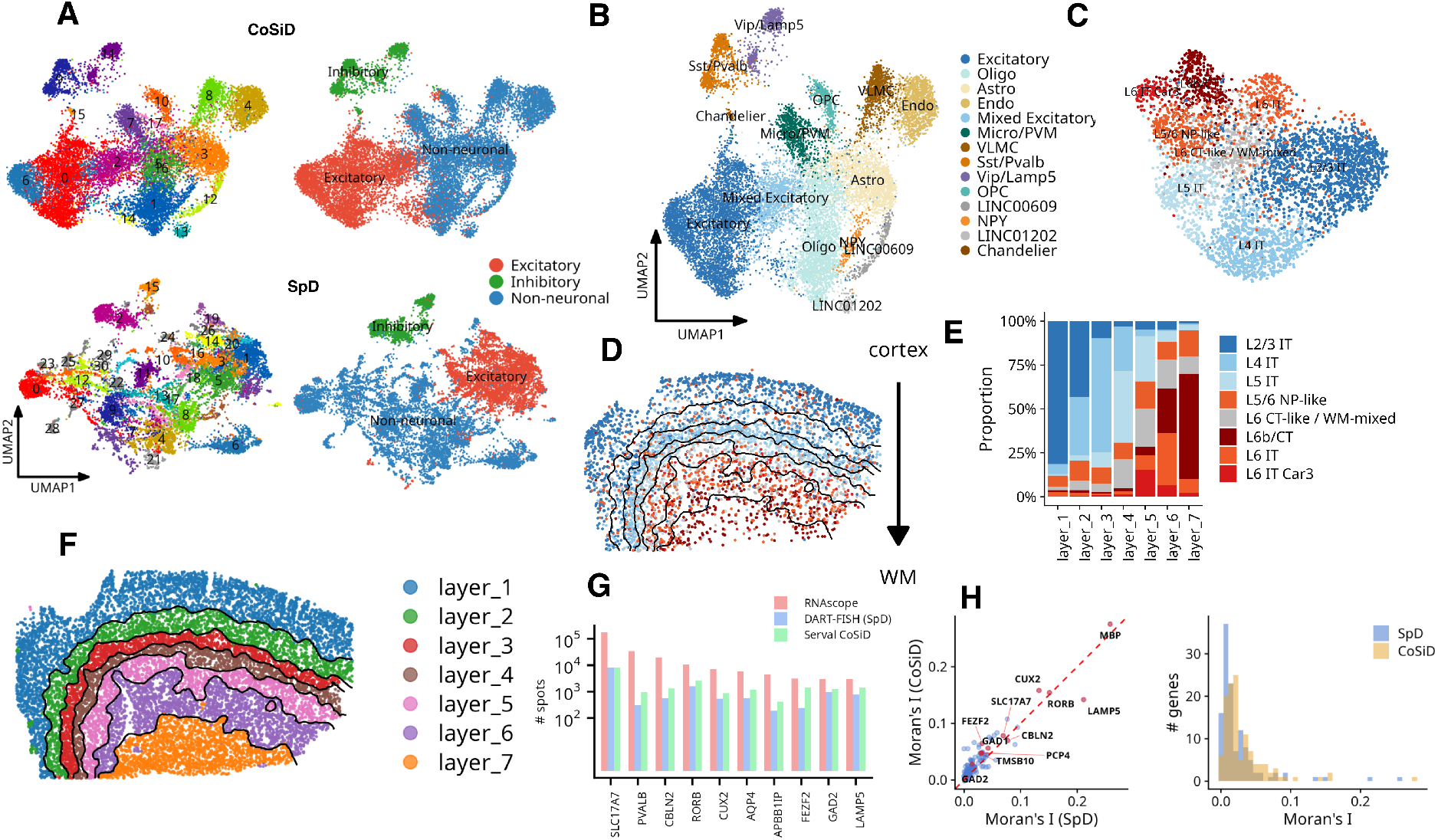
CoSiD generalizes to DART-FISH data and improves cell-type and spatial-domain interpretability. **(A)** UMAP visualization of high-confidence DART-FISH cells decoded with CoSiD (upper) and the original sparse deconvolution decoder (SpD, lower). High-confidence cells were defined as cells receiving concordant labels from scGate marker gating and Seurat label transfer to the snRNA-seq reference. Cells are shown by unsupervised cluster identity and by predicted major class annotation, highlighting improved clustering stability after CoSiD. **(B)** *De novo* annotation of CoSiD-derived clusters in A using marker expression. **(C)** Sub-clustering of CoSiD-decoded excitatory neurons resolves major cortical excitatory subclasses, including L2/3 IT, L4 IT, L5 IT, L6 IT, L6 IT Car3, and deep-layer L6b/CT-like populations. **(D)** Spatial localization of excitatory subclasses across the cortical depth, Cortex to White Matter. **(E)** Composition of SpaMask-inferred layers across the excitatory neurons subset, showing enrichment of upper-layer IT populations in superficial domains and deep-layer subclasses in deeper domains. **(F)** Spatial organization of SpaMask domains across the full CoSiD-decoded dataset. Approximate laminar boundaries were drawn by smoothing the ordered SpaMask layer assignments over a spatial grid using k-nearest-neighbor averaging and plotting contour lines between adjacent inferred layer values. **(G)** Comparison of transcript recovery for selected marker genes between RNAscope, DART-FISH SpD, and CoSiD, reproducing the DART-FISH paper benchmarking analysis. **(H)** Per-gene Moran’s I spatial autocorrelation analysis comparing SpD and CoSiD across the DART-FISH gene panel. Some key cell type markers are highlighted.

The improvement was particularly apparent for the excitatory neuron annotation. The original DART-FISH analysis reported that excitatory Leiden clustering was unstable to cell-order shuffling and therefore required repeated clustering, co-clustering index filtering, centroid correlation to the snRNA-seq reference and manual marker inspection. In contrast, after CoSiD, subclustering of high-confidence excitatory neurons produced marker-coherent subclasses that aligned with expected cortical excitatory populations, including L2/3 IT, L4 IT, L5 IT, L6 IT, L6 IT Car3, and deep L6b/CT-like populations (Figure 4C, Supplementary Figure 13D). Some transitional clusters, such as L5/6 NP-like or L6 CT/white-matter-associated populations, remained less sharply resolved, which is expected given the limited targeted panel and the known difficulty of separating deep-layer subclasses in this dataset [10]. Nevertheless, the CoSiD-derived excitatory clusters were sufficiently structured to support direct annotation using gene markers, laminar position, and the snRNA-seq reference, reducing dependence on iterative cluster-stability filtering and manual curation.

We next quantified the spatial domains by applying SpaMask [15] to recover the expected seven [14] spatial layers (Figure 4E and Figure 4F). The identified excitatory neuronal subclasses colocalized well with the inferred spatial layers, showing progressive changes in excitatory subclass composition across cortical depth (Figure 4D and Figure 4E). Moreover, the inferred domains formed spatially coherent laminar niches across the full tissue (Figure 4F, Supplementary Figure 13C). Using SpaMask with the SpD results, recovered the same broader cortical anatomy (Supplementary Figure 13A). However, the off-diagonal cells reflect reassignment between neighboring layers, indicating differences in inferred boundary placement (Supplementary Figure 13B).

Lastly, to compare with the original DART-FISH benchmarking analysis, we quantified transcript recovery for marker genes previously evaluated against RNAscope in a selected region of interest. CoSiD recovered marker-gene counts higher than what the authors reported for SpD (Figure 4G). Consistent with preservation of spatial structure, per-gene Moran’s I values were broadly concordant between SpD and CoSiD, with CoSiD showing higher overall spatial autocorrelation for most known cortical layer markers (Figure 4H). Thus, CoSiD increased transcript recovery while maintaining and strengthening spatially organized gene-expression patterns.

## Discussion

In this work, we introduce Serval, a modular decoding framework for IST analysis, and demonstrate that it can faithfully reproduce results from the widely used MERlin pipeline for MERFISH decoding [5], while enabling flexible integration of novel decoding algorithms. Building on this framework, we developed CoSiD, which optimizes scaling factors through a principled similarity-based objective. In synthetic benchmarks, CoSiD achieved functional parity with the state-of-the-art MERlin algorithm and outperformed other IST methods including BarDensr [7] and DeepCell-Spots [6]. The analysis of experimental MERFISH data demonstrated that CoSiD improved transcript recovery and bulk RNA-seq correlations while reducing off-target signal and producing cleaner expression matrices for downstream annotation, ultimately improving the biological interpretability of the decoded data. By reprocessing the DART-FISH human primary motor cortex dataset [10], we show that Serval generalizes effectively beyond MERFISH, improving transcript recovery, enhancing cluster resolution, and simplifying reference-guided annotation compared to the original sparse deconvolution approach. These results highlight that the decoding bottleneck is a universal challenge in IST, and Serval provides the infrastructure to address it across platforms.

Prior decoding approaches have typically emphasized heuristic scaling updates (e.g., MERlin) or deep learning-based spot detection (e.g., DeepCell-Spots). While these methods achieve good performance under some conditions, they can be sensitive to imaging variability. By aligning scaling optimization with the decoding objective, CoSiD achieves more stable performance across conditions. Furthermore, Serval’s modular design enables the field to move toward cross-platform benchmarking, providing a standardized environment that is essential as spatial transcriptomics continues to mature.

The practical impact of improved decoding on downstream analysis is seen in the enhanced biological resolution of downstream analyses. CoSiD reduced spurious detection of immune markers in immunodeficient 4T1 NRG tissue, reduced co-expression rate of some mutually exclusive markers (e.g. Neutrophils and Macrophages) and improved clustering coherence of major cell types. In our DART-FISH analysis, the increased sensitivity and improved signal-to-noise ratio simplified the annotation of complex cortical excitatory subclasses, mitigating excessive filtering and reducing the reliance on unstable clustering procedures.

A limitation of our study is the dependence of CoSiD’s performance on appropriate regularization. However, because Serval is modular, these parameters can be refined or replaced as new automated selection algorithms are developed. Finally, while our evaluation focused on MERFISH and DART-FISH, extension to other platforms such as the NanoString CosMx [16] and 10x Xenium [17, 18] will be important for broader adoption. Although these platforms do not currently provide access to the raw spot images, we believe Serval’s architecture establishes a reproducible framework for evaluating decoding strategies, contributing to the establishment of robust standards for high-resolution IST.

## Acknowledgments

Thank you to Eric Lee for providing feedback and advice on generating the figures. 4T1 cells were kindly provided by Gregory Hannon and the IMAXT Laboratory, CRUK Cambridge Institute.

## Declarations

### Author Contributions

J.T. and A.R. conceived the project. J.T., W.C., and L.L. developed the multiplexed MERFISH spot simulator. J.T. and A.R. conceived CoSiD. J.T. generated the synthetic datasets, designed the decoding experiments, and developed CoSiD. J.T. implemented the penalty optimization pipeline and carried out performance benchmarking on synthetic datasets. J.T. implemented and executed the JSIT, DeepCell-Spots, and BarDensr pipelines, including parameter sweeps and comparative evaluations. Y.T. segmented nuclei using CellPose and generated expanded labels to approximate full cell boundaries. N.A. performed downstream analysis of experimental data and wrote the corresponding methods and results sections. S.H. provided guidance on simulator design and microscope setup. D.B., J.M., and X.Z. prepared the C1E1 library and contributed to barcode library design. C.F., S.A., E.K., H.L. and C.O’F. conducted the experiments, generated and processed MERFISH imaging data. Y.T. and I.R. assisted with early software setup for DeepCell-Spots and JSIT, respectively. S.Apa. supervised the wet lab work and oversaw generation of the MERFISH experimental datasets. J.T. benchmarked decoding methods on both synthetic and experimental datasets and performed comparative evaluation. J.T., N.A., and A.R. interpreted the results. J.T. wrote the manuscript with feedback from all authors. A.R. supervised the project.

### Data and Code Availability

All code for the Serval framework and CoSiD are publicly available on GitHub at https://github.com/Roth-Lab/serval. The code for the Multiplexed Spots Simulator is available at https://github.com/Roth-Lab/FISHsim. The pipeline for synthetic experiments is available at https://github.com/Roth-Lab/serval-fishsim-smk. The Serval DART-FISH pipeline is available at https://github.com/Roth-Lab/serval-dartfish-smk. The refactored version for DeepCell-Spots is available at https://github.com/tsuijenk/DeepCell-Spots. The refactored version for JSIT is available at https://github.com/tsuijenk/JSIT. The MERFISH datasets generated in this study have been archived: the 4T1 cell-culture dataset (https://doi.org/10.7910/DVN/NE0WOP) and the 4T1 tissue dataset (https://doi.org/10.5281/zenodo.15678603). The public 4T1 scRNA-seq reference dataset used for MERFISH annotation is available in the NCBI Gene Expression Omnibus (GEO) database under accession code GSE189856 [19].

The DART-FISH human primary motor cortex dataset analyzed in this study is publicly available [10]. Raw registered images and intermediate outputs of the processing pipeline were obtained from Zenodo (https://doi.org/10.5281/ZENODO.8253771). Spot tables, RiboSoma images, and segmentation masks were obtained from Figshare (https://doi.org/10.6084/m9.figshare.23932863.v1) as described by Kalhor et al. (2024). The single-nucleus RNA sequencing reference data for the human primary motor cortex (M1C) from Jorstad et al. (2023) is available for download from the Neuroscience Multi-omics Archive (https://data.nemoarchive.org/publication_release/Human_Cross_Areal_Analysis/).

### Ethics Statement

All animal experimental work was approved by the Animal Care Committee (ACC) and the Animal Welfare and Ethical Review Committee at the University of British Columbia (UBC) under protocol A19-0298.

## References

[1] Ozirmak Lermi, N. et al. Comparison of imaging based single-cell resolution spatial transcriptomics profiling platforms using formalin-fixed paraffin-embedded tumor samples. Nature communications 16, 8499 (2025). URL https://europepmc.org/articles/PMC12474935.

[2] Ya, D. et al. Application of spatial transcriptome technologies to neurological diseases. Frontiers in Cell and Developmental Biology 11 (2023). URL 10.3389/fcell.2023.1142923.

[3] Kang, D. H., Kim, Y., Lee, J. H., Kang, H. S. & Chung, C. Spatial Transcriptomics in lung Cancer and Pulmonary Diseases: A Comprehensive review. Cancers 17, 1912 (2025). URL 10.3390/cancers17121912.

[4] Jin, Y. et al. Advances in spatial transcriptomics and its applications in cancer research. Molecular Cancer 23 (2024). URL 10.1186/s12943-024-02040-9.

[5] Emanuel, G., Eichhorn, S. & Zhuang, X. ZhuangLab/MERlin: MERlin V0.1.6. Zenodo (2020). URL 10.5281/zenodo.3758540.

[6] Laubscher, E. et al. Accurate single-molecule spot detection for image-based spatial transcriptomics with weakly supervised deep learning. Cell Systems 15, 475–482.e6 (2024). URL 10.1016/j.cels.2024.04.006.

[7] Bryan, J. P. et al. Optimization-based decoding of Imaging Spatial Transcriptomics data. Tech. Rep. (2023). URL 10.1093/bioinformatics/btad362.

[8] Chen, S. et al. BARcode DEmixing through Non-negative Spatial Regression (BarDensr). PLoS Computational Biology 17, e1008256 (2021). URL https://pmc.ncbi.nlm.nih.gov/articles/PMC7971881/.

[9] Cisar, C., Keener, N., Ruffalo, M. & Paten, B. A unified pipeline for FISH spatial transcriptomics. Cell Genomics 3, 100384 (2023). URL https://pubmed.ncbi.nlm.nih.gov/37719153/.

[10] Kalhor, K. et al. Mapping human tissues with highly multiplexed RNA in situ hybridization. Nature Communications 15, 2511 (2024). URL 10.1038/s41467-024-46437-y.

[11] Lin, L. A concordance correlation coefficient to evaluate reproducibility. Biometrics 45, 255 (1989). URL 10.2307/2532051.

[12] Demšar, J. Statistical Comparisons of Classifiers over Multiple Data Sets. Tech. Rep. (2006). URL https://www.jmlr.org/papers/volume7/demsar06a/demsar06a.pdf.

[13] Andreatta, M., Berenstein, A. J. & Carmona, S. J. scGate: marker-based purification of cell types from heterogeneous single-cell RNA-seq datasets. Bioinformatics 38, 2642–2644 (2022). URL 10.1093/bioinformatics/btac141.

[14] Jorstad, N. L. et al. Transcriptomic cytoarchitecture reveals principles of human neocortex organization. Science 382, eadf6812 (2023). URL 10.1126/science.adf6812.

[15] Min, W., Fang, D., Chen, J. & Zhang, S. SpaMask: Dual masking graph autoencoder with contrastive learning for spatial transcriptomics. PLoS Computational Biology 21, e1012881 (2025). URL 10.1371/journal.pcbi.1012881.

[16] He, S. et al. High-plex imaging of RNA and proteins at subcellular resolution in fixed tissue by spatial molecular imaging. Nature Biotechnology 40, 1794–1806 (2022). URL https://pubmed.ncbi.nlm.nih.gov/36203011/.

[17] Salas, S. M. et al. Optimizing Xenium In Situ data utility by quality assessment and bestpractice analysis workflows. Nature Methods (2025). URL https://www.nature.com/articles/s41592-025-02617-2.

[18] Janesick, A. et al. High resolution mapping of the tumor microenvironment using integrated singlecell, spatial and in situ analysis. Nature Communications 14 (2023). URL https://www.nature.com/articles/s41467-023-43458-x.

[19] Li, J. et al. Non-cell-autonomous cancer progression from chromosomal instability. Nature 620, 1080–1088 (2023). URL https://www.nature.com/articles/s41586-023-06464-z.

[20] Bindea, G. et al. Spatiotemporal dynamics of intratumoral immune cells reveal the immune landscape in human cancer. Immunity 39, 782–795 (2013). URL 10.1016/j.immuni.2013.10.003.

[21] Newman, A. M. et al. Robust enumeration of cell subsets from tissue expression profiles. Nature Methods 12, 453–457 (2015). URL https://www.nature.com/articles/nmeth.3337.

[22] Rose, S. A. et al. A microRNA expression and regulatory element activity atlas of the mouse immune system. Nature Immunology 22, 914–927 (2021). URL https://www.nature.com/articles/s41590-021-00944-y.

[23] Wagenblast, E. et al. A model of breast cancer heterogeneity reveals vascular mimicry as a driver of metastasis. Nature 520, 358–362 (2015). URL https://www.nature.com/articles/nature14403.

[24] Chen, K. H., Boettiger, A. N., Moffitt, J. R., Wang, S. & Zhuang, X. Spatially resolved, highly multiplexed RNA profiling in single cells. Science 348 (2015). URL https://pubmed.ncbi.nlm.nih.gov/25858977/.

[25] Moffitt, J. R. et al. High-throughput single-cell gene-expression profiling with multiplexed error-robust fluorescence in situ hybridization. Proceedings of the National Academy of Sciences 113, 11046–11051 (2016). URL 10.1073/pnas.1612826113.

[26] Team, C. I. G. C. et al. Dissociation of solid tumor tissues with cold active protease for singlecell RNA-seq minimizes conserved collagenase-associated stress responses. Genome biology 20, 210 (2019). URL 10.1186/s13059-019-1830-0.

[27] Funnell, T. et al. Single-cell genomic variation induced by mutational processes in cancer. Nature 612, 106–115 (2022). URL 10.1038/s41586-022-05249-0.

[28] Rouillard, J.-m. OligoArray 2.0: design of oligonucleotide probes for DNA microarrays using a thermodynamic approach. Nucleic Acids Research 31, 3057–3062 (2003). URL https://pubmed.ncbi.nlm.nih.gov/12799432/.

[29] Xu, Q., Schlabach, M. R., Hannon, G. J. & Elledge, S. J. Design of 240,000 orthogonal 25mer DNA barcode probes. Proceedings of the National Academy of Sciences 106, 2289–2294 (2009). URL https://pubmed.ncbi.nlm.nih.gov/19171886/.

[30] Speiser, A. et al. Deep learning enables fast and dense single-molecule localization with high accuracy. Nature Methods 18, 1082–1090 (2021). URL https://www.nature.com/articles/s41592-021-01236-x.

[31] Babcock, H. P. & Zhuang, X. Analyzing single molecule localization microscopy data using cubic SPlines. Scientific Reports 7 (2017). URL https://pubmed.ncbi.nlm.nih.gov/28373678/.

[32] Stringer, C., Wang, T., Michaelos, M. & Pachitariu, M. Cellpose: a generalist algorithm for cellular segmentation. Nature Methods 18, 100–106 (2020). URL https://www.nature.com/articles/s41592-020-01018-x.

[33] Hao, Y. et al. Dictionary learning for integrative, multimodal and scalable single-cell analysis. Nature Biotechnology 42, 293–304 (2023). URL 10.1038/s41587-023-01767-y.

[34] Amezquita, R. A. et al. Orchestrating single-cell analysis with Bioconductor. Nature Methods 17, 137–145 (2019). URL 10.1038/s41592-019-0654-x.

[35] Plummer, J. T. et al. Standardized metrics for assessment and reproducibility of imaging-based spatial transcriptomics datasets. Nature Biotechnology (2025). URL 10.1038/s41587-025-02811-9.

[36] Lun, A. Clustering algorithms for bioconductor 1.22.0 edn (2026). URL https://mirror.accum.se/mirror/bioconductor.org/packages/3.23/bioc/manuals/bluster/man/bluster.pdf.

[37] Jorstad, N. L. et al. Comparative transcriptomics reveals human-specific cortical features. Science 382, eade9516 (2023). URL 10.1126/science.ade9516.

