## Supplementary Methods and Figures for "Serval: A modular framework for decoding imaging based spatial transcriptomics data"

### Supplemental methods

#### 1 Codebook Library Generation

Both synthetic and experimental datasets used a binary barcode library based on the minimum Hamming distance 4 (MHD4) scheme, a standard in MERFISH experiments. Each barcode consisted of a fixed Hamming weight of 4, ensuring that any two barcodes differ by at least four positions. This encoding supports robust error correction in the presence of missing or noisy fluorescence signals. The codebook spanned 8 hybridization rounds, each imaged in 2 fluorescence channels, resulting in a binary matrix where each row defines a barcode's expected activation pattern across the imaging rounds and channels. Each "1" in the matrix corresponds to a hybridization event designed to be fluorescently labeled and imaged in the specified round and channel. This codebook was shared across all synthetic scenarios and real MERFISH datasets, ensuring consistency in decoding benchmarks. During analysis, the decoding algorithms were provided with the complete codebook as ground truth. Gene targets for the experimental C1E1 MERFISH library were manually curated with inspiration from a few sources: the released Nanostring nCounter gene panels for "Human Pan Cancer Immune Profiling" (converted to mouse homologues), immune gene lists from Bindea et al. [20], suppl. table 1, the leukocyte signature matrix from Newman et al. [21], the marker lists from the transcriptomic data in the Immunological genome project database [22]. In addition, we added genes known to be upregulated in 4T1 cells from RNAseq data obtained by the Hannon laboratory in the course of previous work [23]. The final list of 140 genes was used to design MERFISH probes using the methodology developed by the laboratory of Xiaowei Zhuang as previously described [24, 25].

#### 2 MERFISH Imaging

##### 2.1 Cell culture

4T1 cells were maintained in DMEM supplemented with 10% FBS in a humidified incubator at 37 °C.

##### 2.2 4T1 tumor transplants

4T1 cells were transplanted into the mammary fat pad of 8-week-old NOD/Rag1$^-$/Il2r$^-$ (NRG) mice under anesthesia. Mice were euthanized when the size of the tumors approached 1,000 mm$^3$ in volume, according to the limits of the experimental protocol. Tumor tissue was excised and frozen in optimal cutting temperature (OCT) compound and stored at −80 °C. Tissue was sectioned on a cryostat at −20 °C for downstream preparations.

##### 2.3 Bulk RNA sequencing

Total RNA was extracted from cultured cells or from sections of OCT embedded tissue using Trizol, and RNA integrity was assessed using an Agilent Bioanalyzer. Libraries were then prepped following the standard protocol for the Illumina Stranded mRNA prep (Illumina). Sequencing was performed on the Illumina NextSeq2000 with Paired End 59bp × 59bp reads. Sequencing data was demultiplexed using Illumina's BCL Convert. De-multiplexed read sequences were then aligned to the Mus Musculous (mm10) reference sequence using DRAGEN RNA app on Basespace Sequence Hub. Sequencing was performed at the The School of Biomedical Engineering (SBME) Sequencing Core at University of

British Columbia.

The demultiplexed FASTQ files were then used directly for transcript-level quantification with Kallisto v0.44.0 against the Genome assembly GRCm39 mouse index (NCBI RefSeq assembly: GCF_000001635.27). Each Kallisto run generates an abundance file that, for every transcript, reports its unadjusted length, its effective length (which corrects for Kallisto's fragment-bias model), an estimated fragment count, and a TPM (transcripts per million) value. The effective length is typically slightly shorter than the actual transcript length and reflects the expected mappable region after accounting for biases in where fragments are likely to originate.

TPM itself is calculated in two main steps. First, each transcript's estimated count is divided by its effective length in kilobases to yield a "reads per kilobase" (RPK) value, which corrects for the fact that longer transcripts will attract more reads. Second, all of the RPK values in that sample are summed to produce a single scaling factor, and each transcript's RPK is divided by this factor and multiplied by one million. As a result of that final scaling, the TPM values for every transcript in a sample add up to exactly 1,000,000, so differences in transcript length and sequencing depth are both accounted for, and the abundances can be compared directly across samples.

In the final bulk file, TPMs for all transcript isoforms were summed per gene, version suffixes were stripped from Ensembl IDs, and gene symbols were appended, yielding a gene-level TPM matrix for subsequent analysis.

### 3 Single cell RNA sequencing

For single cell RNA sequencing, cultured 4T1 cells were trypsinized, pelleted and resuspended in 0.04% BSA/PBS. 4T1 tumour fragments were dissociated into single cells with cold active Bacillus lichenformis (Creative Enzymes NATE0633) in PBS supplemented with 5 mM CaCl2 and 125 Uml-1 DNAse, as described previously [26, 27]. Cells were then pelleted and resuspended in 0.04% BSA/PBS and immediately loaded onto a 10X Genomics Chromium single-cell controller targeting 3,000 cells for recovery. Libraries were prepared according to the 10X Genomics Single Cell 3' Reagent kit standard protocol. Libraries were then sequenced on an Illumina Nextseq500/550 at the School of Biomedical Engineering (SMBE) Sequencing Core at University of British Columbia, or a HiSeqX at the Michael Smith's Genome Sciences Centre targeting 50,000 reads per cell. 10X Genomics Cell Ranger 3.0.2 was used to perform demultiplexing, counting and alignment to mm10.

#### 3.1 Sample Preparation

Cultured 4T1 cells were fixed in 3.2% paraformaldehyde (PFA) in phosphate-buffered saline (PBS) for 15 minutes, followed by permeabilization with 0.5% Triton X-100 for 10 minutes at room temperature. 4T1 tumor tissue sections were fixed in 3.2% paraformaldehyde (PFA) in phosphate-buffered saline (PBS) for 15 minutes and were then transferred to 70% ethanol and stored overnight at 4 °C.

#### 3.2 Hybridization and Gel Embedding

Prior to hybridization, samples were equilibrated in a formamide-containing buffer and incubated with encoding probes in a hybridization mixture containing yeast tRNA, dextran sulfate, and mruine RNase inhibitors. After hybridization, samples were embedded in polyacrylamide and enzymatically cleared using a proteinase K-based digestion buffer. Cleared gels were stored in 2× saline-sodium citrate (SSC) buffer in RNase inhibitors and ProClin at 4 °C until imaging.

##### 3.3 Imaging Hardware and Acquisition

Samples were mounted in a Bioptech's FCS2 flow chamber for fluid handling. Imaging was performed on a Nikon ECLIPSE Ti2-E inverted microscope equipped with a 60× oil-immersion objective lens (NA = 1.4) and a Photometrics Prime 95B sCMOS camera. Excitation was delivered via a BrixXHUB laser combiner with four laser lines: 473 nm, 561 nm, 647 nm, and 750 nm. Laser intensities were set to 2%, 14%, 20%, and 30% power, respectively. Emission was collected through a quad-band dichroic filter set (ZET405/473/561/647–656/752m; Chroma Technology, batches 326652 and 326205).

##### 3.4 Fluorescence Acquisition

Each sample underwent eight hybridization-imaging rounds with readout probes 5'-conjugated with either ATTO565 or Cy5 and an internal Thiol C6-SS modification to allow cleavage with TCEP during imaging rounds. Two bits per round were imaged using either 561 nm or 647 nm excitation. Z-stacks were collected with 1 $\mu$m step size to cover the full focal volume. Exposure times were 0.5 s (473 nm, fiducial beads), 1 s (561 nm, target and Syto85 nuclear stain), 0.75 s (647 nm), and 1 s (750 nm).

##### 3.5 Autofocus and Synchronization

Autofocus was achieved via a two-step procedure: first, a coarse z-stack was evaluated using Brenner's gradient focus metric; second, fiducial beads were segmented across z-planes and best focus was refined using multi-scale extrema detection. Hardware-based transistor–transistor logic (TTL) triggering synchronized camera, laser, and stage operations. Exposure output was set to `"Any Row"` or `"All Rows"` to align with the laser triggering logic. Excitation followed the order: 473 nm, 647 nm, 750 nm, 561 nm. Rolling shutter synchronization was validated using oscilloscope traces.

##### 3.6 Fluidics System and Fluorophore Cleaving

Between imaging rounds, fluorophores were cleaved using tris(2-carboxyethyl)phosphine (TCEP), delivered by a custom-built fluidics system. This system utilized programmable syringe pumps (30 mL for general buffer flow and 10 mL dedicated to TCEP) and three manually actuated multi-valve ports (MVP1–3) to route reagents through eight fluid channels. Valve configurations minimized dead volume and isolated corrosive reagents. A MATLAB-based control script orchestrated fluidics, XY stage, z-drive, laser shutters, and camera via hardware triggers.

##### 3.7 Probe Design

Probes were designed as 90-nt oligonucleotides comprising, from 5' to 3', a 20-nt forward (index) primer, a 30-nt readout sequence, a 30-nt target-binding region, a second 30-nt readout sequence, and a 40-nt reverse (index) primer (including a T7 promoter) [24]. To maximize hybridization efficiency, target-binding regions were drawn from the most abundant transcript isoform of each gene (Gencode v18) and filtered in OligoArray2.1 to yield 30-nt sequences with melting temperature $> 70$ °C, no predicted secondary structure above 76 °C, and no runs of $\geq 6$ identical nucleotides [28]. Candidate probes were further vetted by BLAST + against the mouse transcriptome to eliminate off-target matches.

Index primers were selected from a library of orthogonal 25-mers [29] trimmed to 20-nt, each with a 3' GC clamp and melting temperature of 70–80 °C. A T7 promoter (`TAATACGACTCACTATAGGG`) was appended to the reverse primer to enable in vitro transcription. Readout sequences (30-nt) were constructed by concatenating 20-nt primer motifs with 10-nt segments of distinct primer sequences; each was screened by BLAST + to avoid complementarity with index primers, target regions, or the mouse genome.

Probes were assembled in silico by concatenating index primers, readout sequences, and target-binding regions such that all possible barcode–readout pairings were uniformly represented and randomly distributed along each transcript. The final set of $\sim$ 100,000 oligonucleotides was synthesized as an array-derived complex pool (Twist Biosciences) and amplified with index primers to generate the working MERFISH library.

Readout probes were conjugated with a 5' Cy5 or ATTO565 and a Thiol S-S modification to facilitate cleavage with TCEP. Fluorescent probes were from Bio-Synthesis or Biomers.net.

##### 3.8 Customized Fluidics and Optical Path Enable Reliable MERFISH Imaging

To enhance imaging consistency and minimize system noise, we implemented several custom hardware modifications. A bespoke syringe pump design ensured stable fluid exchange while eliminating pulsatile fluctuations. A dedicated reagent line for TCEP cleavage was included to prevent cross-contamination. In contrast to default vendor laser configurations, our system employed high-power laser lines for 561 nm and 647 nm excitation. Additionally, Köhler illumination was adopted to ensure even excitation across the imaging field.

#### 4 MERFISH Datasets

We evaluated decoding performance on both real MERFISH datasets and synthetically generated multiplexed images.

The *cultured cells dataset* consists of MERFISH images acquired from 4T1 cell cultures, a murine breast cancer cell line grown under controlled in vitro conditions. The dataset includes 15 fields of view (FOVs), each with 15 z-slices, acquired at a spatial resolution of 0.024 $\mu$m per pixel. Each FOV covers a $1608 \times 1608$ pixel region.

The *tissue dataset* was derived from 4T1 tumors implanted in mouse tissue, representing an in vivo biological environment with increased complexity in background fluorescence and tissue structure. This dataset contains 54 FOVs with 28 z-slices per field and a spatial resolution of 0.174129 $\mu$m per pixel. Each FOV spans $1608 \times 1608$ pixels. These two datasets together span a range of experimental complexity, allowing evaluation of decoding performance under both controlled and biologically heterogeneous conditions.

In addition to real data, we generated a suite of synthetic datasets using our custom MERFISH image simulator (see The Image Simulator: Camera Model and Noise Types section). Each synthetic image was constructed using known barcoded spot identities and subjected to physical and optical noise consistent with experimental MERFISH. Synthetic datasets varied in signal-to-background properties, photon count per barcode, and round-to-round intensity variation, enabling controlled benchmarking across a range of decoding difficulty.

#### 5 Multiplexed Spot Simulation

We developed a physically grounded simulator to generate synthetic MERFISH images that reflect the optical, physical, and noise characteristics of real imaging experiments. This simulation framework is inspired by the general image formation pipeline of DECODE [30], but tailored specifically for multiplexed barcoded imaging. Our simulator includes realistic models for fluorescent blinking, photon emission, camera noise, and background variation, enabling precise benchmarking of decoding algorithms under controlled yet photophysically grounded conditions.

The simulator generates one image per hybridization round and imaging channel, incorporating a diffraction-limited point spread function (PSF) to model optical blur. Each fluorescent emitter is modeled as a point object and convolved with a 3D PSF, estimated via cubic spline interpolation. We use the following convolutional model:

$$I(\vec{r}) = O(\vec{r}) \circledast PSF(\vec{r}) \tag{1}$$

where $O(\vec{r})$ is the object signal, $PSF(\vec{r})$ is the spatially calibrated point spread function, and $\circledast$ denotes convolution. The PSF is modeled as a spline-weighted sum following the form described by Babcock et al. [31], with 64 coefficients per voxel and per z-slice.

We incorporate a range of noise sources observed in experimental MERFISH data, including:

- **Photon shot noise**: modeled as a Poisson process with quantum efficiency (QE) scaling per wavelength.
- **Dark current noise**: thermal signal accumulation, suppressed via cooling assumptions.
- **Readout noise**: amplifier-dependent, non-Gaussian per-pixel variation modeled using RMS statistics.
- **Quantization noise**: rounding errors from analog-to-digital conversion based on a 16-bit ADC.

The simulator includes background modeling in two forms: (i) *cell-local background*, reflecting autofluorescence or cytoplasmic haze, controlled via a signal-to-cell-background ratio (SCR); and (ii) *global background*, generated via a kernel mixture model to introduce randomized spatial heterogeneity. Additional parameters simulate barcode dropout and extra-bit false positives, based on empirical observations from real MERFISH images. This simulation framework serves both to evaluate decoder robustness under varying optical conditions and to benchmark against known ground truth spot locations and identities. Details for multiplexed spot simulator can be found in Supplementary Note 1: Multiplexed Spot Simulator Details. Details for the various simulation scenarios can be found in Supplementary Table 1.

#### 6 Decoding Methods

##### *MERlin*

The original MERlin pipeline implements a heuristic decoding procedure that combines fiducial-based registration, deconvolution, and signal normalization. Barcode assignment is performed by minimizing the Euclidean distance between intensity vectors of candidate pixels and known barcode signatures. This distance is computed after vector normalization and per-round scaling, which implicitly acts like a heuristic Expectation-Maximization (EM) procedure, though this is not explicitly stated.

##### *Serval Implementation of MERlin*

To improve transparency and modularity, we developed Serval Merlin (Figure 1), a refactored version of the MERlin pipeline that decouples each processing stage. Serval Merlin retains the core decoding steps, starting from chromatic correction, followed by fiducial alignment, image filtering (high-pass, deconvolution, low-pass), per-round scaling factor initialization, decoding, and barcode filtering. This design enables more consistent benchmarking across decoders and reduces entanglement of preprocessing with decoding logic.

**Supplementary Table 1**: Summary of simulation scenarios used to evaluate decoder performance. Scenarios 1–10 constitute the primary benchmark.

| Scenario | Photon Level | SCR | Background | Bit Dropout | Bit Addition | Description |
|---|---|---|---|---|---|---|
| 1 | High, variable | High (4.0) | 0.05 | 0 | 0 | Baseline simulation |
| 2 | Moderate | Moderate (1.2) | 0.08 | 0 | 0 | Reduced signal-to-cell contrast |
| 3 | Low | Moderate (1.0) | 0.10 | 0 | 0 | Low-photon imaging condition |
| 4 | Low | Very low (0.2) | 0.10 | 0 | 0 | Severe low-contrast condition |
| 5 | Uniform low | Moderate (1.2) | 0.08 | 0 | 0 | Uniform signal intensity across rounds |
| 6 | High, variable | Moderate-high (1.5) | 0.05 | 0.08 | 0 | Barcode dropout errors |
| 7 | High, variable | Moderate-high (1.5) | 0.05 | 0.15 | 0 | Severe barcode dropout errors |
| 8 | High, variable | Moderate-high (1.5) | 0.05 | 0 | 0.05 | Barcode addition errors |
| 9 | High, variable | Variable (0.7–1.8) | 0.10 | 0.03 | 0.05 | Mixed realistic degradation |
| 10 | Low, variable | Variable (0.5–1.5) | 0.25 | 0.05 | 0.08 | Combined degradation benchmark |

##### *Cosine Optimized Pixel Decoder*

We propose CoSiD as a decoding improvement over the Serval Merlin baseline. Rather than relying on Euclidean distance, CoSiD maximizes the cosine similarity between pixel intensity vectors (across imaging rounds) and barcode signatures. This approach emphasizes the angular similarity of a signal trace while discounting global intensity magnitude. It is particularly robust to variation in signal strength across rounds. To enhance stability, CoSiD includes two regularization strategies: (i) it penalizes scaling factors that deviate too far from 1.0, and (ii) it promotes scaling entropy to prevent overfitting to dominant bits. This balance improves sensitivity and pattern robustness while maintaining decoding accuracy. See Supplementary Note 2: CoSiD Optimization for full objective formulation and gradient derivation.

##### *Simple Nearest Neighbor Decoder*

As a baseline, we implemented a Simple Nearest Neighbor decoder. It assigns barcode identities based on the closest match between pixel vectors and codebook entries using Euclidean distance. To ensure robustness, pixels with low norm or poor similarity are filtered out using thresholding. This approach lacks scaling normalization and serves as a minimal, model-free comparator.

##### *Benchmark Baselines*

For comparative evaluation, we also benchmarked three recent decoding pipelines: JSIT [7], DeepCell-Spots [6], and BarDensr [8]. JSIT is an optimization-based decoder that reconstructs the observed image as a sparse combination of codebook signatures convolved with a point spread function (PSF). DeepCell-Spots applies a weakly supervised convolutional neural network trained to predict spot probabilities from input images. BarDensr uses a non-negative matrix factorization (NMF) model to estimate spatial barcode densities by modeling the image as a sum of barcode templates weighted by local abundance. For all three methods, we performed parameter sweeps over key hyperparameters to ensure fair comparisons. Default configurations were used as baselines where applicable. See Supplementary Note 3: Practical Guidance for Selecting Regularization Parameters for detailed sweep settings.

##### *Pipeline Evolution*

Our decoding pipeline evolved from the original MERlin to CoSiD through a modular transition. We reimplemented the core functionality of MERlin to support controlled comparison, then modularized its components to form Serval Merlin. The decoding logic was subsequently replaced with CoSiD. This progressive transformation enables reproducible benchmarking and targeted performance evaluation.

#### 7 Evaluation of MERlin and Serval Merlin

In order to assess the equivalence of MERlin and Serval Merlin, we compared their decoded outputs at the single-cell level. Specifically, we evaluated the total transcript counts per cell between the two methods and reported Spearman correlation coefficient along with its associated $p$-value, deeming the correlation statistically significant when $p < 0.001$. To further quantify absolute agreement, we computed Lin's Concordance Correlation Coefficient [11], a standard metric for evaluating agreement with a reference method. To visualize distributional similarities, we also generated side-by-side violin plots of the transcript counts per cell from both methods. Finally, we constructed a Bland-Altman Plot, where the $x$-axis represents the mean transcript counts per cell between the two methods, and the $y$-axis represents the difference in transcript counts per cell between the methods. This visualization allowed us to assess potential biases or systematic deviations between methods across the dynamic range of transcript counts.

#### 8 Decoder Benchmarking and Metric Computation on Synthetic Datasets

To benchmark decoder performance, synthetic MERFISH images were generated using our multiplexed spot simulator across ten primary scenarios, with 100 replicates per scenario. The scenarios systematically varied photon counts, signal-to-cell-background ratios (SCR), barcode bit drop probability, barcode bit add probability, background sampling probability, round-specific photon variation, and complete round failure.

Decoder performance was compared among CoSiD, the Serval implementation of MERlin, a Nearest Neighbor baseline, BarDensr, and DeepCell-Spots. Because the simulated images were generated using a fixed point spread function (PSF), all ground-truth transcripts occupied a single pixel and no area filtering was applied during evaluation.

For each decoder, performance was evaluated across the full range of decoder-specific scores. CoSiD, MERlin, and Nearest Neighbor were evaluated using barcode mean distance, whereas BarDensr and DeepCell-Spots were evaluated using evidence and probability scores, respectively. Because these distinct decoders produce continuous outputs on different numerical scales, applying a single, fixed threshold would introduce evaluation bias. To ensure equitable comparison, transcript-level and abundance-recovery metrics were systematically evaluated across a sweep of all score thresholds for each unique simulation run, replicate, and decoder.

Transcript-level detection performance was assessed using exact recall, exact false discovery rate ($FDR$), exact $F_1$ score, exact average precision ($AP$), localization recall, and localization $F_1$ score. Exact metrics required both correct transcript identity and localization, whereas localization metrics considered spatial matching irrespective of transcript identity. Spatial matches were defined using a nearest-neighbor tolerance of one pixel between decoded and ground-truth transcript coordinates. From the raw true positive ($TP$), false positive ($FP$), and false negative ($FN$) counts captured across the score sweep, metrics were dynamically computed. The final operating threshold for each individual replicate and evaluation mode was chosen by identifying the threshold that maximized the local $F_1$ score. All reported point-specific transcript metrics reflect values captured at this optimal $F_1$ operating point. Additionally, to decouple our primary benchmarks from single-threshold selection dependencies, we integrated across the continuous precision-recall curves to compute the Average Precision ($AP$).

Abundance recovery was assessed using Pearson and Spearman correlation coefficients between decoded and ground-truth transcript abundances. For each replicate, Pearson and Spearman correlation coefficients were computed across all score thresholds. The maximum Pearson correlation coefficient and maximum Spearman correlation coefficient obtained across the threshold range were used for downstream comparisons.

Statistical comparisons among decoders were performed using Friedman tests followed by post-hoc Nemenyi tests, with Bonferroni correction applied to control the family-wise error rate. Results are reported as mean $\pm$ standard deviation across the ten primary simulation scenarios unless otherwise stated.

#### 9 Segmentation method of nuclear-stained channel

In our MERFISH experiments, segmentation relied exclusively on the nuclear-stained channel to define cell regions, with each nucleus mask serving as the primary label for spot assignment. After decoding barcodes into spot coordinates and gene identities, these nuclear masks were expanded outward to approximate whole-cell boundaries, and each transcript was assigned the unique identifier of its enclosing nucleus.

All three-dimensional image stacks were first registered to correct lateral offsets and stage drift between imaging rounds. MERlin's fiducial-correlation warp function (high-pass sigma = 3, default settings) was applied to each FOV so that decoded spots and segmentation masks occupied the same spatial frame. From each registered FOV, the single z-slice exhibiting optimal focus and highest signal-to-noise ratio was selected for segmentation.

Nuclear segmentation was carried out using Cellpose [32]. For the cell culture dataset, where nuclear morphology remained essentially constant along the z-axis, the mask generated on that single best-focus slice was propagated across the entire z-stack. In tissue samples, where cell shape and density change with depth, propagation was constrained to an appropriate number of adjacent slices in which each nucleus was expected to appear. For example, an image of 1608 $\times$ 1608 pixels covering 180 $\mu$m $\times$ 280 $\mu$m yields 0.1741 $\mu$m per pixel; an average nucleus of 75 pixel diameter ($\approx$ 13 µm) would span on the order of ten to fifteen 1-$\mu$m z-steps, and so masks were applied across that range of neighbors before excluding more distant slices. Segmentation parameters were set to balance detection sensitivity and specificity: a flow threshold of 0.8 and a cell probability threshold of 0.8. For cell culture images, manual segmentation was performed via Cellpose's graphical interface, and the model was iteratively trained on these user-defined masks to improve its performance. For tissue images, an initial pre-trained nuclei model from our Cambridge collaborators served as the starting point and was progressively refined through human-in-the-loop corrections.

Whole-cell regions were approximated by expanding nuclear seeds outward. A binary "cell mask" was generated by thresholding the decoded spot image to capture each cell's overall silhouette. Nuclear labels then served as seeds for a watershed algorithm that propagated boundaries until adjacent seeds or the cell-mask perimeter were reached, yielding contiguous masks approximating both nuclear and cytoplasmic extents without a dedicated cytoplasmic stain.

Decoded spots were overlaid onto the segmentation masks, and each spot's (x, y) coordinate was mapped to the underlying region to assign it to a specific cell. Regions at the periphery of fields of view were excluded on an experiment-by-experiment basis to remove artifacts arising from imaging overlap and border issues. The resulting, cell-labeled and filtered spot dataset was then exported for downstream gene-per-cell quantification.

#### 10 Performance Evaluation on MERFISH Experimental Datasets

We evaluated decoding performance on two MERFISH experimental datasets: a cultured cell line and a tissue section from a transplanted 4T1 tumor. Unlike synthetic datasets with known ground truth, experimental validation relied on concordance with orthogonal transcriptomic measurements and agreement with the established decoding standard: MERlin.

As in the synthetic analysis, we modulated each method's detection stringency by sweeping the post-decoding parameter that governs transcript recovery, tracing the relationship between transcript recovery and reference correlation rather than collapsing to a single operating point. For the distance-based decoders (CoSiD, Nearest Neighbor, Serval MERlin, and MERlin), which produce connected-component spot detections, we applied a fixed minimum spot area of 3 pixels and swept over a set of mean distance values. JSIT likewise yields connected-component spots, each with an associated area and intensity; for JSIT we first selected a minimum spot area and a minimum signal intensity threshold by grid search, then swept a quantile cutoff on the per-spot decoded signal strength of the retained spots. BarDensr and DeepCell-Spots do not produce connected-component spots and have no analogous area filter; for these we swept the native confidence parameter exposed by each: the spot-calling threshold for BarDensr, and the spot-detection probability and gene-assignment probability for DeepCell-Spots. The full sweep was retained to characterize the transcript-recovery–correlation relationship for each method (Figure 3A, Supplementary Figure 10E).

##### 10.1 Global Expression Concordance

We computed Pearson and Spearman correlation coefficients between pseudo-bulk gene counts (decoded outputs) and matched bulk RNA-seq TPM values. These were visualized as correlation-versus-transcript count plots to reveal trade-offs between sensitivity and concordance.

##### 10.2 Cell-Resolved Transcriptomic Profiling.

We compared decoded outputs from CoSiD and MERlin at the cellular level. Across all segmented cells, we computed:

- total transcript counts per cell,
- unique gene counts per cell,
- total transcript counts per unit cell area, and
- CoSiD–MERlin difference in transcript counts per field of view.

To evaluate whether these differences were statistically significant, we applied the paired Wilcoxon signed-rank test to each metric, comparing CoSiD and MERlin across matched cells. In all cases, the differences were significant if $p < 0.001$. To quantify the magnitude of these differences, we computed Cohen's $d$ effect sizes to reveal the effect of each metric.

#### 11 Serval DART-FISH decoding

We implemented a Snakemake workflow to process the published DART-FISH data [10]. As the supplied images were already registered, no fiducial alignment was performed. The remaining decoding steps including chromatic correction, high pass filtering, deconvolution and low pass filtering where done the same way as the MERFISH workflow. The original SpD based decoding performed maximum projection across z-stacks. We found that this performed poorly for CoSiD, so we instead independently decoded every fourth z-slice. Spots from all decoded z-slices within FOV were combined, and then deduplicated based on minimum distance of 5.5 pixels for spots assigned to the same barcode. Subsequent filtering was done as described below for the original DART-FISH SpD results.

#### 12 DART-FISH data preprocessing, quality control and dimensionality reduction

Two transcript-to-cell assignment strategies were generated: a nuclear-restricted assignment (0-pixel distance) and a permissive 150-pixel radial expansion ($\approx 42.6\ \mu$m, given the 284 nm brain pixel size) for initial comparisons of assignment radius effects. Note that these were regenerated from the DART-FISH segmentation pipeline, as the published brain cell-by-gene matrix was not deposited; the 150-pixel value reflects the *max_rol2nuc_dist* parameter setting [10]. Matrices were imported into Seurat (v5.0.3) [33] and cell metadata (area, centroid coordinates and labels) were obtained from output deposited by the authors. Cells with fewer than 5 total counts were removed.

For quality control, an adaptive upper bound on total counts was defined as the median plus three median absolute deviations (MADs) of the total count distribution per condition to remove extreme outliers. The percentage of negative-control probe counts (probes matching "blank" for CoSiD or "empty" for

DART-FISH) was computed per cell, analogous to mitochondrial-content quality control. Cells were then removed if total counts exceeded the 3 MADs, if control-probe content exceeded 10% of total counts, and if segmented cell area exceeded 6,000. Counts were then log-normalized, reduced by Seurat's RunPCA, and a shared-nearest-neighbor graph was constructed on the first 30 principal components. Cells were then clustered with the Louvain algorithm (0.6 resolution). Seurat's Uniform Manifold Approximation and Projection (UMAP) was computed on the same 30 principal components for visualization.

#### 13 MERFISH data preprocessing, quality control and dimensionality reduction

MERFISH data preprocessing followed the same pipeline as DART-FISH with the following modifications. Cells with fewer than 10 total counts were removed. The upper bound on total counts was defined as the median plus three MADs per condition. Cells were additionally removed if segmented cell area exceeded 15,000, or if the percentage of negative-control probe counts was identified as an upper 3 MADs outlier on the log scale. Counts were log-normalized using scuttle (1.12.0) [34]. Dimensionality reduction and UMAP were computed on the first 40 principal components. For the 4T1 cell culture dataset, clustering at resolution 0.5 identified a low-quality cluster characterised by low total counts, which was removed prior to downstream analysis. The 4T1 scRNA-seq dataset was similarly processed.

#### 14 Spatial Quality Control (QC) Metrics

SpatialQM [35] was adopted for all the spatial QC metrics reported in the downstream comparative analyses. In SpatialQM framework, sparsity was computed as the fraction of zero entries in the count matrix (*coop::sparsity*, R v0.6.3).Higher values indicate higher proportion of zero entries and lower values indicate a denser matrix. Entropy was computed as the Shannon entropy (base 2) of the pooled count values across the expression matrix via the *BioQC::entropy* function. Cluster separation was assessed using approximate silhouette width computed via the *approxSilhouette* function from the bluster R package (v1.12.0) [36]. PCA embeddings were used as the input coordinate space for all silhouette calculations.

#### 15 Annotating MERFISH dataset

Cells were annotated based on known marker genes and reference expression data from a publicly available 4T1 scRNA-seq dataset [19]. Cell-type signatures were derived by identifying differentially expressed genes using the MERFISH panel genes on the public scRNA-seq data. Those were then used to construct a gating model using scGate (v1.6.0) [13]. Cancer cells were identified by positivity for Padi4, Gata3, and Akt3, with negativity for the pan-immune marker Ptprc and the myeloid markers Csf1r and Csf3r. Neutrophils were gated on Csf3r positivity, with negativity for the macrophage markers Csf1r and Msr1. Macrophages were identified by positivity for Msr1 and Csf1r, with negativity for the neutrophil marker Csf3r and the endothelial marker Adamts7. Endothelial cells were defined by positivity for Adamts7 and Scarf1, with negativity for Ptprc. Cells that did not pass any gate or received conflicting assignments were retained as an Unknown or Multi class, respectively.

#### 16 Annotating the DART-FISH Brain dataset

Major cell classes were assigned by two independent approaches and reconciled to retain high-confidence labels. Seurat's reference-based label transfer was performed against a snRNA-seq reference of human primary motor cortex (M1C) [37]. Transfer anchors were identified between reference and query by projection onto the reference principal-component space, and reference Class labels were transferred across these anchors using the first 30 dimensions. In parallel, marker-based gating was performed with scGate (v1.6.0) [13] using signatures for excitatory (positive: SLC17A7, SATB2; negative: GAD2) and inhibitory (positive: GAD1, GAD2; negative: SATB2) neurons; cells assigned to neither gate were designated non-neuronal. Cells receiving concordant labels were then designated as high confidence and subsequently reprocessed and clustered at 0.6 resolution for subclasses *de novo* annotation. Seurat's FindAllMarkers was used to find differentially expressed marker genes (Wilcoxon rank-sum test). Only CoSiD output was annotated. A second round of clustering for the excitatory subset was performed where Louvain resolution was varied until 0.9 to capture the 7 layer subclasses.

#### 17 Spatial domain identification and evaluation

Spatial domains were identified with SpaMask [15], a dual-masking graph autoencoder with contrastive learning. For each condition, the normalized expression matrix and cell centroid coordinates were provided as input, and a spatial neighborhood graph was constructed by k-nearest neighbors ($k = 21$). The model was trained for 500 epochs (hidden dimension 512, latent dimension 256, node feature-mask rate 0.5, edge-drop rate 0.2, contrastive-loss weight $\lambda = 2$), and the latent representation was partitioned into 7 spatial domains by k-means clustering.

### Supplementary Note 1: Multiplexed Spot Simulator Details

#### The Image Simulator: Point Spread Function (PSF)

A PSF represents the response of a microscope to a point source. In the MERFISH context, a point source is a single emitter that forms an image in each of the 16 imaging rounds. Each emitter $O(\vec{r})$ is placed on an object plane and convolved with the PSF to produce an image $I(\vec{r})$:

$$I(\vec{r}) = O(\vec{r}) \circledast PSF(\vec{r}) \tag{2}$$

where $\circledast$ denotes the convolution operator. We use cubic spline functions to approximate the PSF, capturing its shape more accurately than Gaussian fitting. Following Babcock et al.[31], the 3D PSF is modeled as:

$$F_{i,j,k}(x,y,z) = \sum_{m=0}^{3}\sum_{n=0}^{3}\sum_{p=0}^{3} a_{i,j,k,m,n,p} \left(\frac{x-x_i}{dx}\right)^m \left(\frac{y-y_j}{dy}\right)^n \left(\frac{z-z_k}{dz}\right)^p \tag{3}$$

where $dx$, $dy$, and $dz$ are the voxel dimensions; $x_i$, $y_j$, $z_k$ are the voxel coordinates; and $a_{i,j,k,m,n,p}$ are the spline coefficients (64 per voxel). The simulator supports subpixel localization by using continuous-valued emitter coordinates.

#### The Image Simulator: Cell Background Context

MERFISH images exhibit visible cell background autofluorescence, which impacts spot visibility. We control this using the signal-to-cell-background ratio (SCR):

$$SCR = \frac{\text{signal}}{\text{cell-background}} \tag{4}$$

In our empirical datasets, SCR values typically ranged from 1.10 to 1.60 across both bright and dim signals. This range reflects the contrast levels observed in real MERFISH images and modulates how prominently a spot appears against its local cellular background.

#### The Image Simulator: Global Background Context

Experimental MERFISH images also show spatially varying background due to scattered radiance or illumination heterogeneity. We simulate this using a Parzen window kernel method, drawing background values from a mixture of Gaussian distributions to create a randomized spatial profile. This adds complexity and realism by eliminating prior assumptions on background structure.

#### The Image Simulator: Camera Model and Noise Types

We simulate both sample-related and camera-related noise based on sCMOS characteristics. Each layer is detailed below:

##### Photon Shot Noise

Caused by the stochastic nature of photon arrival. The expected detected electron count per pixel $k$ is:

$$\lambda_k = \lambda_{0,k} \cdot qe \tag{5}$$

where $\lambda_{0,k}$ is the photon count and $qe$ is the wavelength-dependent quantum efficiency. The observed signal $s_k$ follows a Poisson distribution:

$$p_{\text{shot}}(s_k) = \frac{\lambda_k^{s_k} e^{-\lambda_k}}{s_k!} \tag{6}$$

Photon shot noise is a physical limit and cannot be eliminated.

##### Dark Current Noise

Light-independent noise from thermally induced electrons. It accumulates over time and is temperature-dependent. Cooling the sensor by every 7°C typically halves the dark current. In our simulator, this is added as a Poisson-distributed offset per pixel.

##### Readout Noise

Noise introduced during signal amplification. CCD/EMCCD sensors exhibit Gaussian-distributed readout noise. In sCMOS sensors, each pixel has a dedicated amplifier, resulting in pixel-dependent and skewed noise distributions. We model both RMS and median noise:

- **RMS noise**: captures full variance including outliers.

- **Median noise**: insensitive to extremes, represents central tendency.

##### Quantization Noise

Arises during analog-to-digital conversion. The quantization step equivalence (QSE) converts electrons to digital counts:

$$QSE = \frac{N_{\text{well}}}{N_{\text{DR}}} \quad \text{(electrons/count)} \tag{7}$$

where $N_{\text{well}}$ is the full well capacity and $N_{\text{DR}}$ is the ADC's dynamic range. We use a 16-bit ADC, yielding $2^{16} = 65,536$ levels.

#### Barcode-Level Perturbations

To simulate realistic decoding errors, we introduce barcode-level dropout and false positives. Each spot has a small probability of dropping one bit (false negative) or acquiring an extra bit (false positive), consistent with artifacts observed in real data.

#### Summary

This simulator models spatial optics, biological variability, and camera physics to generate synthetic MERFISH images under multiple noise conditions. It supports benchmarking decoding pipelines with precise ground truth annotations and simulates emitter placement with subpixel resolution.

### Supplementary Note 2: CoSiD Optimization

#### Objective Function

As outlined in the main text, CoSiD assigns barcodes to candidate pixels by maximizing the cosine similarity between pixel intensity vectors and codebook barcodes. To ensure stable scaling factors across imaging rounds, we optimize a penalized objective function, $\mathcal{L}_p(\mathbf{c})$. Let $\mathbf{c} = [c_1, c_2, \ldots, c_d]$ represent the scaling vector. Conditioned on fixed barcode assignments $\mathbf{b}_i$, the objective is:

$$\mathcal{L}_p(\mathbf{c}|\mathbf{b}) = \sum_{i=1}^{N} \langle \hat{\mathbf{x}}_i(\mathbf{c}), \mathbf{b}_i \rangle - \alpha N \sum_{j=1}^{d} (c_j - 1)^2 - \beta N \sum_{j=1}^{d} \bar{c}_j \log \bar{c}_j \tag{8}$$

where:

- $\mathbf{x}_i$ represents the pre-scaled intensity vector (denoted as $\tilde{\mathbf{x}}_i$ in the main text)
- $\hat{\mathbf{x}}_i(\mathbf{c}) = \dfrac{(\mathbf{x}_i/\mathbf{c})}{\|(\mathbf{x}_i/\mathbf{c})\|_2}$ is the normalized scaled intensity vector for pixel $i$
- $\mathbf{b}_i$ is the matched barcode vector from the codebook, namely

$$\mathbf{b}_i(\mathbf{c}) = \arg\max_{\mathbf{b} \in \mathcal{B}} \langle \hat{\mathbf{x}}_i(\mathbf{c}), \mathbf{b} \rangle$$

- $N$ is the number of decoded pixels used for optimization
- $\alpha$ is the regularization coefficient for the L2 penalty
- $\beta$ is the regularization coefficient for entropy
- $\bar{c}_j = \dfrac{c_j}{\sum_k c_k}$ is the normalized scaling weight per round

During each scaling update step, the barcode assignments $\mathbf{b}_i$ obtained from the current decoder parameters are held fixed while optimizing $\mathbf{c}$.

#### Gradient Derivation

The gradient $\nabla \mathcal{L}_p(\mathbf{c}|\mathbf{b}(\mathbf{c}))$ is computed as follows. Let $\mathbf{y}_i = \mathbf{x}_i/\mathbf{c}$ and define $\hat{\mathbf{y}}_i = \mathbf{y}_i/\|\mathbf{y}_i\|_2$. Then:

$$\frac{\partial \mathcal{L}_p}{\partial c_j} = \sum_{i=1}^{N} \left[ -\frac{1}{\|\mathbf{y}_i\|_2} \left( \mathbf{b}_i^j - (\hat{\mathbf{y}}_i \cdot \mathbf{b}_i) \cdot \hat{\mathbf{y}}_i^j \right) \cdot \frac{x_i^j}{c_j^2} \right] - 2\alpha N(c_j - 1) - \beta N \left( \frac{1}{Z} \log \bar{c}_j + \frac{1}{Z} \right) \quad (9)$$

where $Z = \sum_k c_k$ is the normalization constant. This expression accounts for all three contributions: cosine alignment, L2 stability, and entropy regularization, appropriately scaled by $N$ to remain invariant to batch size. The gradients are optimized using `scipy.optimize.minimize` with positivity constraints on $c_j > 0$.

Although the barcode assignments $\mathbf{b}_i$ depend on the scaling factors $\mathbf{c}$ through the decoding step, we treat such assignments as fixed during the optimization of $\mathbf{c}$. Under this conditional objective, the gradient expression above is exact. After updating $\mathbf{c}$, decoding is rerun on the next batch of images, producing updated barcode assignments.

#### Optimization Procedure

We initialize all scaling factors to 1 and perform bounded optimization using the L-BFGS-B method, a quasi-Newton optimization algorithm that approximates the inverse Hessian matrix in a memory-efficient manner. L-BFGS-B is specifically designed for high-dimensional problems with simple box constraints. In our case, we enforce positivity on all scaling values by setting bounds $c_j > 0$. The method iteratively updates the scaling vector using the analytically derived gradient and maintains a limited memory of past updates to efficiently approximate second-order curvature information.

During each scaling update step, barcode assignments are first obtained by decoding the current batch of images using the current scaling factors. Pixels belonging to valid connected components are collected and grouped by their assigned barcode. The scaling factors are then optimized while holding these assignments fixed, corresponding to maximizing the conditional objective $\mathcal{L}_p(\mathbf{c}|\mathbf{b}(\mathbf{c}))$. After updating the scaling factors, decoding is re-run on subsequent images batches, producing updated barcode assignments. This procedure corresponds to a block coordinate optimization (EM-like) scheme performed over image batches.

Because the number of candidate pixels can be extremely large, training is performed using batches of images rather than the entire dataset simultaneously. The objective and gradient are computed jointly over all selected pixels within each batch. The optimized scaling factors are then applied globally across the dataset prior to final decoding, ensuring stable and consistent barcode assignment under diverse imaging conditions.

### Supplementary Note 3: Practical Guidance for Selecting Regularization Parameters

CoSiD introduces two regularization coefficients: an L2 penalty ($\alpha$) anchoring scaling factors to 1 and an entropy term ($\beta$) promoting diversity across imaging rounds. To evaluate parameter sensitivity, we performed systematic hyperparameter analyses on both a MERFISH cell-line dataset and a MERFISH tissue dataset. Parameter sweeps were conducted across a broad range of entropy and L2 penalties, and decoding performance was assessed using correlation to matched bulk RNA-seq measurements, blank-barcode detection rates, and preservation of expected biological structure.

For the MERFISH cell-line dataset, the strongest performance was obtained for parameter combinations near ($\alpha = 0.01, \beta = 2.0$), while the MERFISH tissue dataset achieved its best performance near ($\alpha = 0.002, \beta = 1.0$). These analyses demonstrated that decoder performance exhibits dataset-dependent behavior and that quantitative agreement with bulk RNA-seq measurements can be improved through targeted hyperparameter optimization. In contrast, the default parameter values ($\alpha = 0.001, \beta = 0.01$) were used without modification for both the synthetic and DART-FISH datasets and produced satisfactory decoding performance. Consequently, we recommend the default parameters as a general starting point, with additional tuning reserved for applications requiring maximal quantitative agreement with external abundance measurements or when systematic artifacts are observed during quality-control assessment.

As practical diagnostics, we recommend examining (i) correlation to bulk RNA-seq measurements, (ii) blank-barcode detection rates, and (iii) preservation of expected marker-gene structure in downstream analyses. Substantial deviations in these quantities may indicate under- or over-regularization and can guide selection of appropriate parameter values for new datasets.

### Supplementary Note 4: Parameter Sweep for Benchmark Baselines

#### BarDensr

BarDensr applies non-negative matrix factorization to estimate spatial barcode densities from multiplexed images. In our benchmarking, we used the standard preprocessing steps of min-max normalization followed by background subtraction, consistent with the authors' guidelines. The only tunable parameter was the `spot_calling_threshold`, which controls the confidence level required to report a decoded spot. We performed a grid sweep over the following values:

- `spot_calling_threshold` = {0.25, 0.30, 0.35, 0.40, 0.45, 0.50, 0.55, 0.60, 0.65, 0.70, 0.75, 0.80}

All other parameters were fixed at their default values.

#### DeepCell-Spots

DeepCell-Spots applies a two-stage decoding process. During first stage, it uses a weakly supervised convolutional neural network to predict spot probabilities per imaging round, classifying each pixel into foreground or background. Subsequently, for each barcoded vector of probability values, we fit a graphical model of a mixture of relaxed Bernoulli distributions using stochastic variational inference.

The only tunable decoding parameter was the `spot_detection_probability_threshold` in the first stage, which determines the minimum detection probability required for a region to be classified as a spot in each imaging round. This threshold is passed into the Polaris model and directly controls the sensitivity of spot calling.

- `spot_detection_probability_threshold` = {0.05, 0.1, 0.2, 0.4, 0.6, 0.8, 0.95}

These values span lenient (0.05, 0.1, 0.2), moderate (0.4, 0.6), and stringent (0.8, 0.95) spot acceptance. For each of these spot detection probability threshold, we plotted their correlation results against bulk-seq at each assignment probability threshold (0, 0.2, 0.4, 0.6, 0.8, 0.95, 0.99), which is an output from assigning gene identities in second stage.

We were unable to run the original author's source code since they were calling a version of gene assignment function that no longer existed; hence, we made modifications to their source code and created a refactored version that has the capability to run in parallel across FOV/Z on compute clusters. GitHub link to the refactored version can be found in Data and Code Availability.

#### JSIT

JSIT reconstructs the observed image using sparse representations of codebook signatures convolved with a PSF. Among the tunable parameters are the proximal operator type along with its regularization strength and hyperparameter. We tested their default proximal operator, SGL, but it returned minimal decoded barcodes and pronounced undercalling, whereas the $L0$ operator yielded correlation and detected-transcript counts comparable to the other decoding methods. We therefore used the $L0$ operator with $\lambda = 5$ and $\beta = 200$. We applied JSIT's default PSF width $\sigma = 1.25$ and image scaling factor $sf = 3$ to both datasets.

JSIT produces connected-component spot detections, reporting for each decoded barcode its area (number of pixels), its mean signal intensity in the observed image, and a decoded signal strength, defined as its mean intensity in JSIT's reconstructed signal map; both intensities are averaged over the pixels of the barcode's connected component. To trace the transcript-recovery–correlation relationship for JSIT, we applied a two-stage post-decoding sweep. We first performed a grid search over minimum spot area and minimum signal intensity thresholds, retaining the combination that maximized reference correlation. On the retained barcodes, we then swept a quantile cutoff on the per-spot decoded signal strength, from the 0th to the 90th percentile in steps of 10, progressively removing the lowest intensity detections to traverse the transcript-recovery axis. At each quantile we recomputed the correlation between decoded pseudo-bulk abundances and the bulk reference, yielding the JSIT curve in Figure 3A and Supplementary Figure 10E. The GitHub link to the refactored version can be found in Data and Code Availability.

### Supplemental figures and tables

**Supplementary Table 2**: Performance summary across synthetic benchmarking scenarios. Values are reported as mean ± standard deviation across all simulation replicates. Exact metrics require correct transcript identity and localization, whereas localization metrics evaluate spatial recovery independent of transcript identity. Pearson $r$ and Spearman $\rho$ quantify abundance recovery relative to ground-truth transcript abundances.

| Decoder | Exact Recall | Exact FDR | Exact F1 | Pearson $r$ | Spearman $\rho$ | Exact AP | Localization F1 | Localization Recall |
|---|---|---|---|---|---|---|---|---|
| CoSiD | $0.864 \pm 0.033$ | $0.051 \pm 0.006$ | $0.904 \pm 0.020$ | $0.975 \pm 0.024$ | $0.972 \pm 0.023$ | $0.968 \pm 0.015$ | $0.906 \pm 0.020$ | $0.868 \pm 0.033$ |
| MERlin | $0.868 \pm 0.032$ | $0.050 \pm 0.005$ | $0.907 \pm 0.019$ | $0.978 \pm 0.014$ | $0.975 \pm 0.016$ | $0.967 \pm 0.021$ | $0.909 \pm 0.019$ | $0.872 \pm 0.032$ |
| BarDensr | $0.712 \pm 0.182$ | $0.230 \pm 0.214$ | $0.739 \pm 0.198$ | $0.528 \pm 0.226$ | $0.558 \pm 0.221$ | $0.773 \pm 0.244$ | $0.848 \pm 0.174$ | $0.901 \pm 0.116$ |
| Nearest Neighbor | $0.492 \pm 0.158$ | $0.185 \pm 0.051$ | $0.606 \pm 0.127$ | $0.270 \pm 0.248$ | $0.344 \pm 0.241$ | $0.790 \pm 0.100$ | $0.618 \pm 0.123$ | $0.503 \pm 0.155$ |
| DeepCell-Spots | $0.634 \pm 0.217$ | $0.259 \pm 0.249$ | $0.679 \pm 0.232$ | $0.267 \pm 0.449$ | $0.722 \pm 0.256$ | $0.733 \pm 0.247$ | $0.743 \pm 0.253$ | $0.740 \pm 0.250$ |

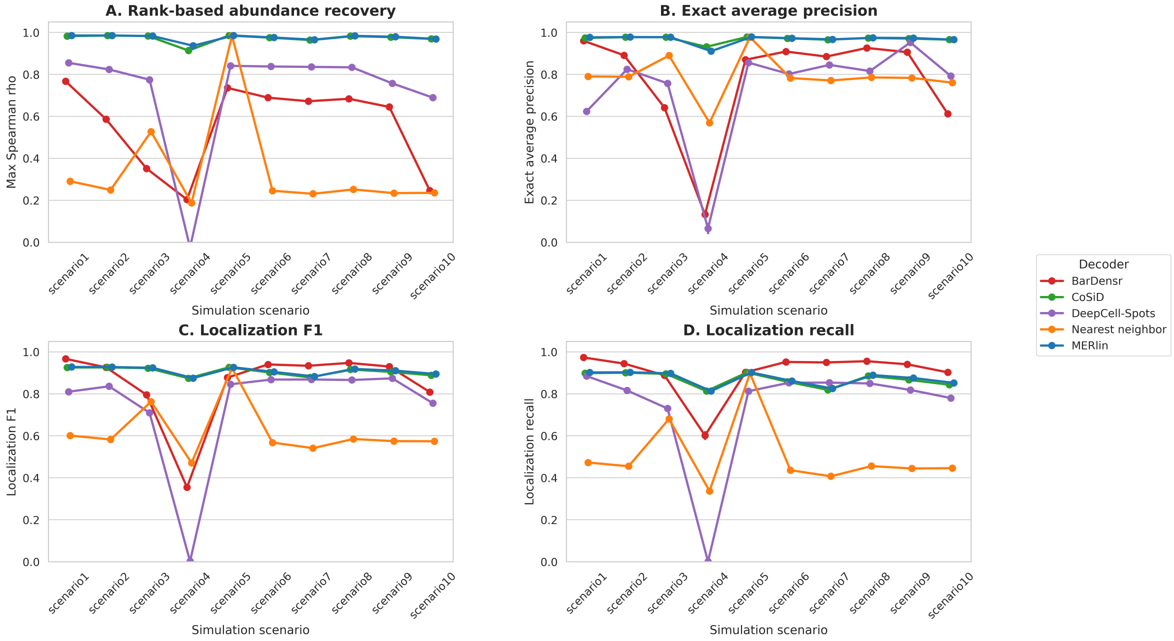


**Supplemental Figure 1**: **CoSiD matches MERlin performance and exceeds competing decoding methods in synthetic benchmarks.** **(A)** Maximum Spearman correlation coefficient between decoded and ground-truth transcript abundances across the ten primary simulation scenarios defined in Supplementary Table 1. **(B)** Exact average precision (AP). **(C)** Localization F1 score. **(D)** Localization recall. Points represent the mean across 100 replicates per scenario and error bars indicate the standard error of the mean. Lines connect each decoder across scenarios to aid visual tracking of performance and do not imply continuity or ordering between scenarios. Decoder performance is shown for CoSiD (orange), MERlin (purple), BarDensr (blue), Nearest Neighbor (red), and DeepCell-Spots (green).

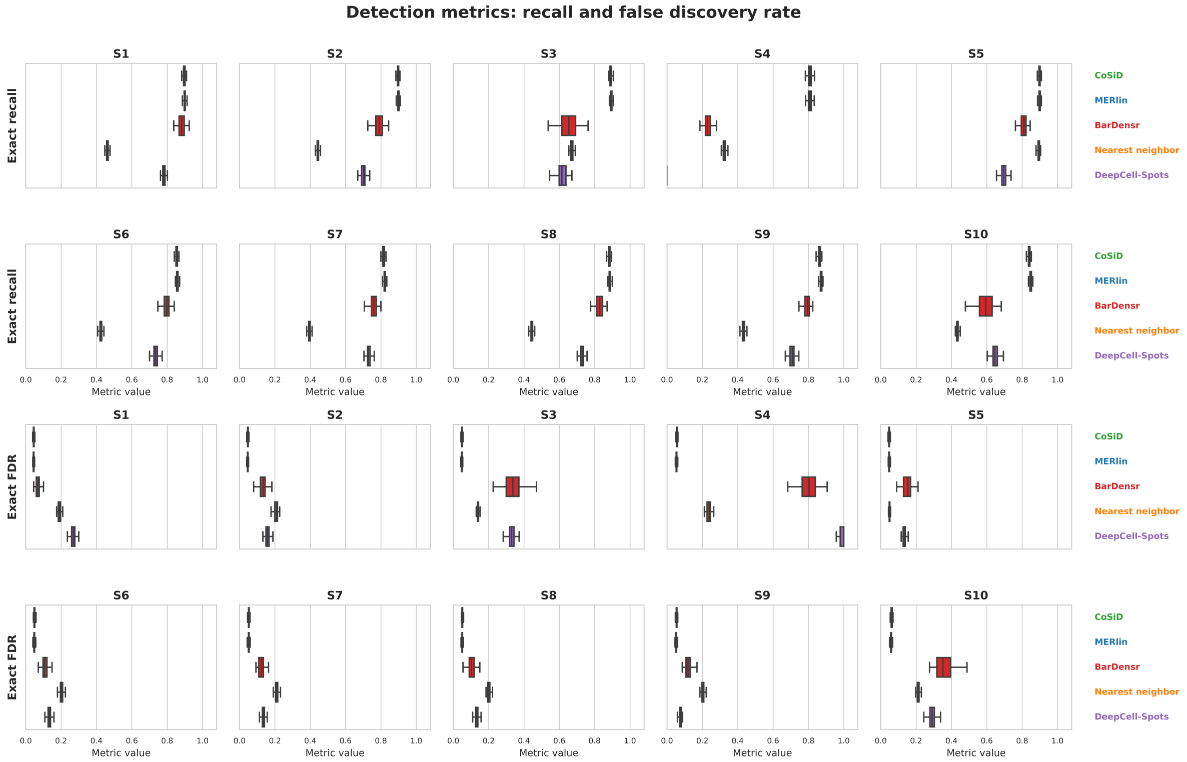


**Supplemental Figure 2**: **Per-scenario distributions of transcript-level detection accuracy: exact recall and false discovery rate.** Exact recall (top two rows) and exact false discovery rate (FDR; bottom two rows) for each decoder across the ten synthetic benchmarking scenarios (S1–S10, arranged as two rows of five; 100 replicates per scenario; scenario definitions in Supplementary Table 1). Decoders (CoSiD, MERlin, BarDensr, Nearest neighbor, DeepCell-Spots) are coloured consistently with Figure 1. Boxes show the interquartile range (IQR) and median; whiskers extend to 1.5×IQR; outliers are omitted. Note that for exact FDR, *lower* values indicate better performance, opposite to all other metrics in these supplementary figures, for which higher values are better.

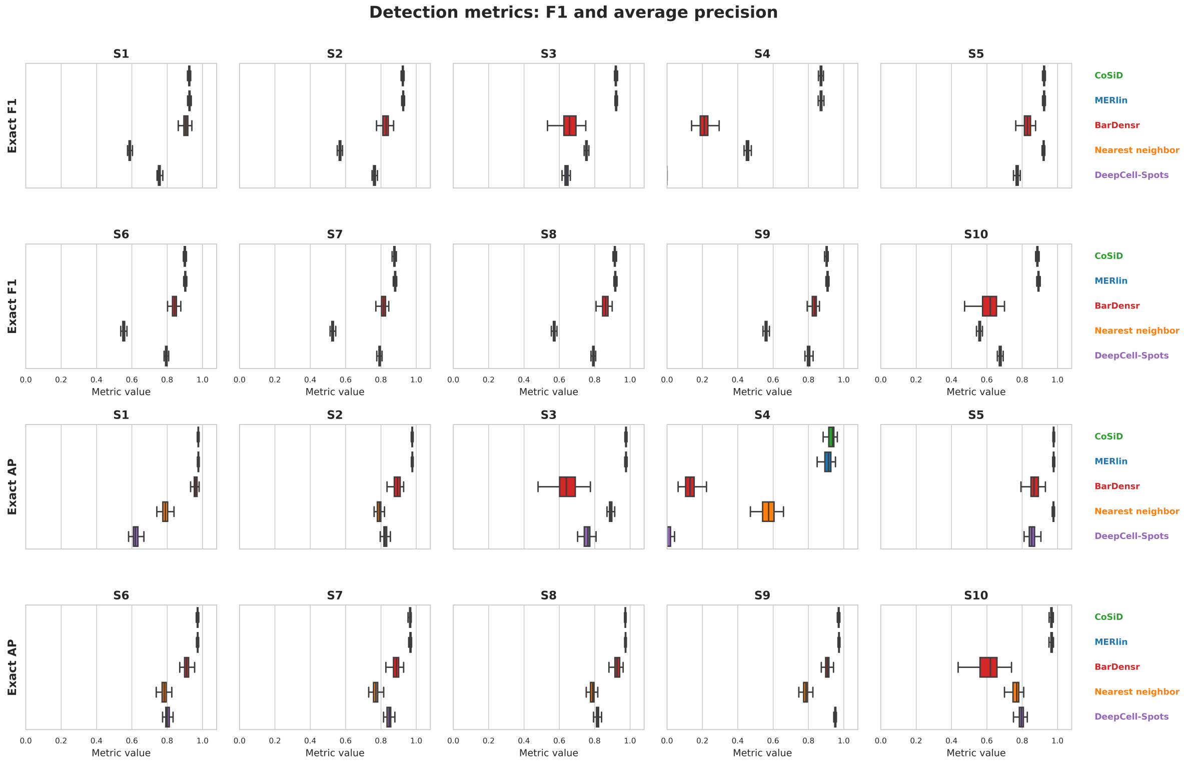


**Supplemental Figure 3**: **Per-scenario distributions of transcript-level detection accuracy: F1 score and average precision.** Exact F1 score (top two rows) and exact average precision (AP; bottom two rows) for each decoder across the ten synthetic benchmarking scenarios (S1–S10, arranged as two rows of five; 100 replicates per scenario; scenario definitions in Supplementary Table 1). Decoders (CoSiD, MERlin, BarDensr, Nearest neighbor, DeepCell-Spots) are coloured consistently with Figure 1. Boxes show the interquartile range (IQR) and median; whiskers extend to 1.5×IQR; outliers are omitted. For both metrics, higher values indicate better performance.

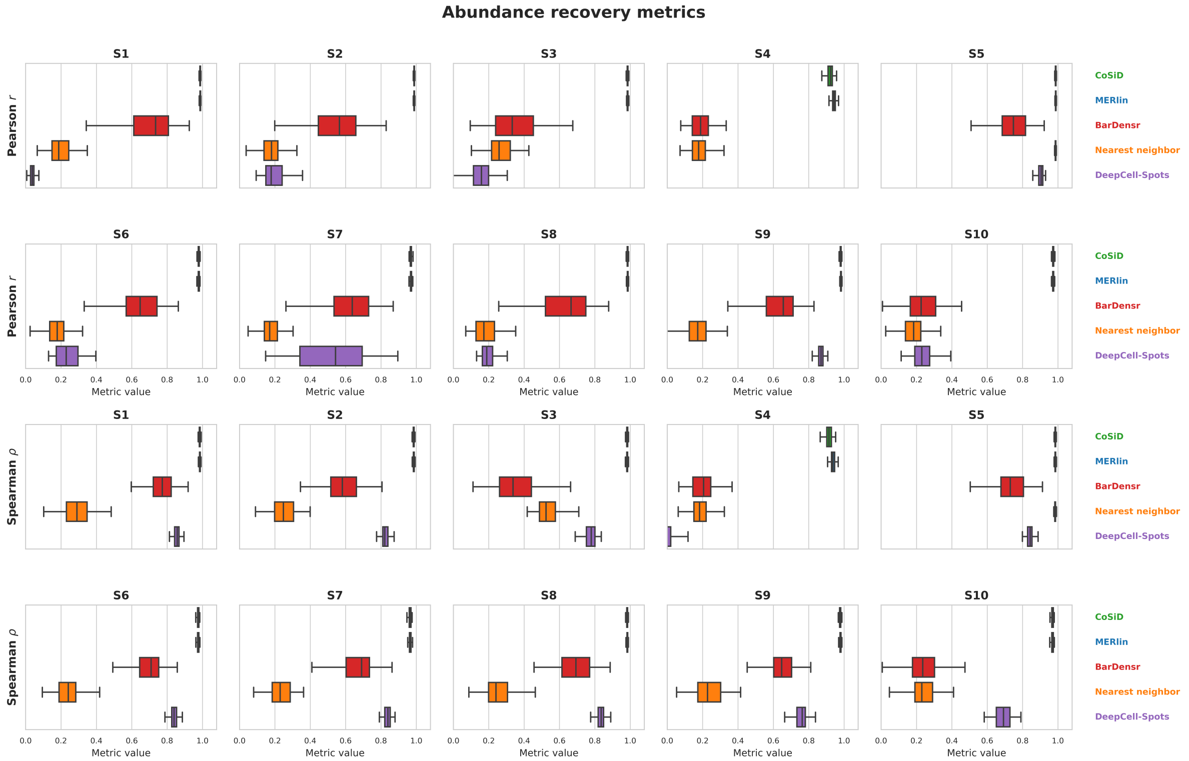


**Supplemental Figure 4**: **Per-scenario distributions of abundance recovery.** Maximum Pearson correlation coefficient ($r$; top two rows) and maximum Spearman correlation coefficient ($\rho$; bottom two rows) between decoded and ground-truth transcript abundances, for each decoder across the ten synthetic benchmarking scenarios (S1–S10, arranged as two rows of five; 100 replicates per scenario; scenario definitions in Supplementary Table 1). Decoders (CoSiD, MERlin, BarDensr, Nearest neighbor, DeepCell-Spots) are coloured consistently with Figure 1. Boxes show the interquartile range (IQR) and median; whiskers extend to 1.5×IQR; outliers are omitted. For both metrics, higher values indicate better performance.

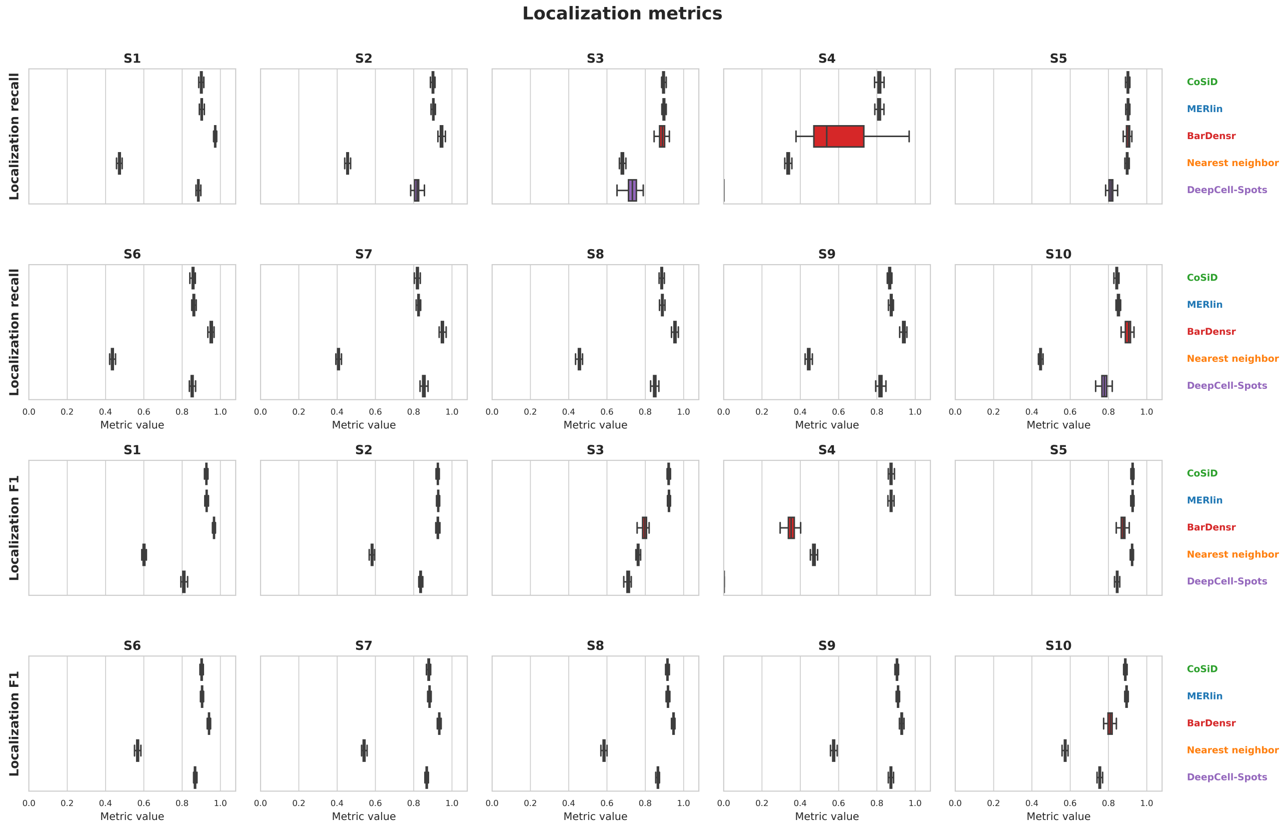


**Supplemental Figure 5**: **Per-scenario distributions of localization accuracy.** Localization recall (top two rows) and localization F1 (bottom two rows), which assess spatial recovery of detected spots independent of transcript identity, for each decoder across the ten synthetic benchmarking scenarios (S1–S10, arranged as two rows of five; 100 replicates per scenario; scenario definitions in Supplementary Table 1). Decoders (CoSiD, MERlin, BarDensr, Nearest neighbor, DeepCell-Spots) are coloured consistently with Figure 1. Boxes show the interquartile range (IQR) and median; whiskers extend to 1.5×IQR; outliers are omitted. For both metrics, higher values indicate better performance.

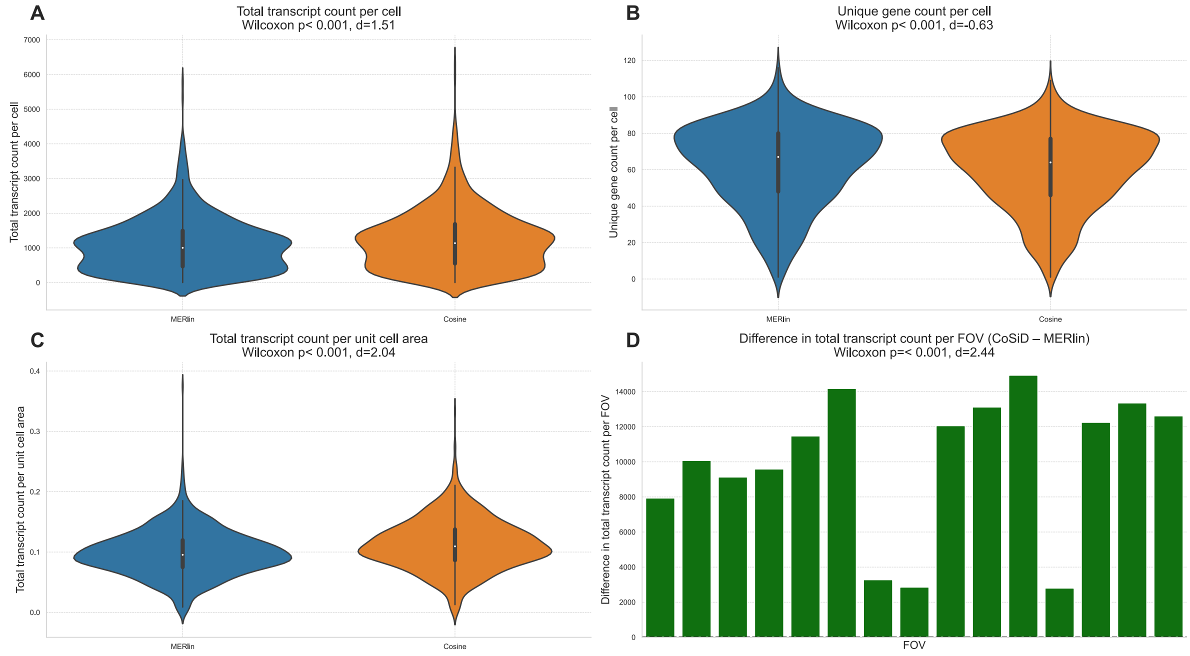


**Supplemental Figure 6**: **Comparison of MERlin and CoSiD on 4T1 cell-line MERFISH data.** **(A)** Violin plots of total transcript counts per cell for MERlin (blue) and CoSiD (orange) across 1017 cells (Wilcoxon signed-rank $p < 0.001$, Cohen's $d = 1.51$). **(B)** Violin plots of unique gene counts per cell ($p < 0.001$, $d = -0.63$). **(C)** Violin plots of total transcript counts normalized by cell area ($p < 0.001$, $d = 2.04$). In **(A-C)**, violins show the full distribution, central bars denote median $\pm$ interquartile range. **(D)** Bar plot of per-FOV differences in total transcript recovery (CoSiD - MERlin) across 15 fields of view ($p < 0.001$, $d = 2.44$). CoSiD yields consistently higher transcript counts at both the single-cell and FOV-level.

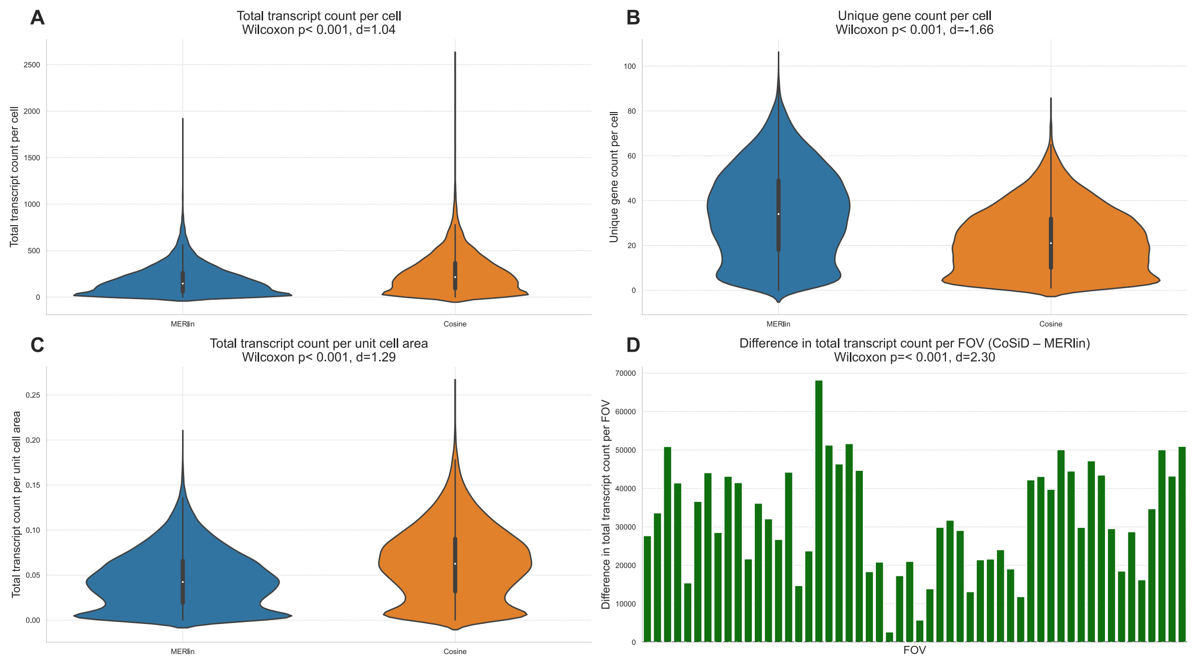


**Supplemental Figure 7**: **Comparison of MERlin and CoSiD on 4T1 tissue MERFISH data.** **(A)** Violin plots of total transcript counts per cell for MERlin (blue) and CoSiD (orange) across 22705 cells (Wilcoxon signed-rank $p < 0.001$, Cohen's $d = 1.04$). **(B)** Violin plots of unique gene counts per cell ($p < 0.001$, $d = -1.66$). **(C)** Violin plots of total transcript counts normalized by cell area ($p < 0.001$, $d = 1.29$). In **(A-C)**, violins show the full distribution, central bars denote median $\pm$ interquartile range. **(D)** Bar plot of per-FOV differences in total transcript recovery (CoSiD - MERlin) across 54 fields of view ($p < 0.001$, $d = 2.30$). CoSiD yields consistently higher transcript counts at both the single-cell and FOV-level.

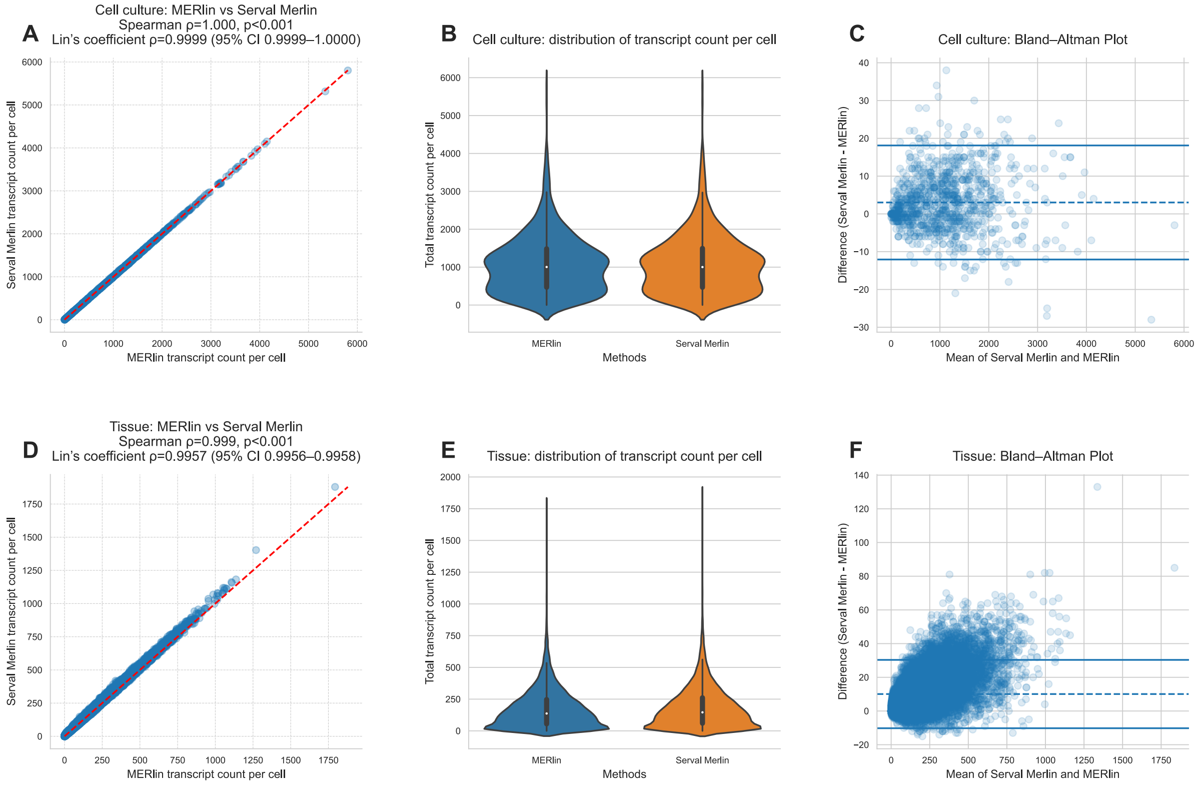


**Supplemental Figure 8**: **Comparison of MERlin and Serval Merlin decoding on cell-line and tissue MERFISH data. (A)** Scatterplot of per-cell transcript counts in the 4T1 cell-line dataset, showing near-perfect concordance between MERlin and Serval Merlin (Spearman's $\rho = 1.000$, $p < 0.001$; Lin's concordance $\rho = 0.9999$, 95% CI 0.9999–1.0000). **(B)** Violin plots of the same cell-line transcript distributions for MERlin (blue) and Serval Merlin (orange). **(C)** Bland–Altman analysis for the cell-line data; dashed line denotes mean bias, solid lines denote the 95% limits of agreement. **(D)** Scatterplot of per-cell counts in the tissue dataset (Spearman's $\rho = 0.999$, $p < 0.001$; Lin's concordance $\rho = 0.9957$, 95% CI 0.9956-0.9958). **(E)** Violin plots of transcript distributions per cell in tissue. **(F)** Bland–Altman plot for the tissue data.

**Supplementary Table 3**: **Standardized spatial quality control metrics for MERFISH 4T1 tissue.** Comparison of overall transcript recovery, density, and matrix sparsity between CoSiD and the default MERlin decoder.

| Decoder | Panel Size | N Cells | Transcripts/Cell | Transcripts/Area | Sparsity | Entropy |
|---|---|---|---|---|---|---|
| CoSiD | 134 | 20,765 | 280.220 | 0.070 | 0.819 | 1.170 |
| MERlin | 134 | 20,765 | 197.569 | 0.049 | 0.723 | 1.665 |

Metrics computed using SpatialQM framework. CoSiD vs MERlin default decoder.

**Supplementary Table 4**: **Silhouette width clustering evaluation by cell type.** Assessment of cluster separation quality across annotated cell types in the 4T1 dataset. CoSiD yields higher median silhouette widths and significantly lower fractions of cells with negative silhouette scores (particularly in Cancer cells and Neutrophils), indicating tighter and more distinct transcriptional clusters compared to MERlin.

| | CoSiD | | | MERlin | | |
|---|---|---|---|---|---|---|
| | **N** | **Median sil.** | **Frac. negative** | **N** | **Median sil.** | **Frac. negative** |
| **Cancer.cell** | 5,297 | 0.030 | 20.2% | 7,231 | −0.009 | 59.2% |
| **Endothelial** | 503 | 0.134 | 2.4% | 208 | 0.156 | 3.4% |
| **Macrophage** | 3,794 | 0.027 | 28.3% | 3,338 | 0.022 | 35.1% |
| **Neutrophil** | 256 | 0.137 | 3.9% | 358 | 0.069 | 20.9% |

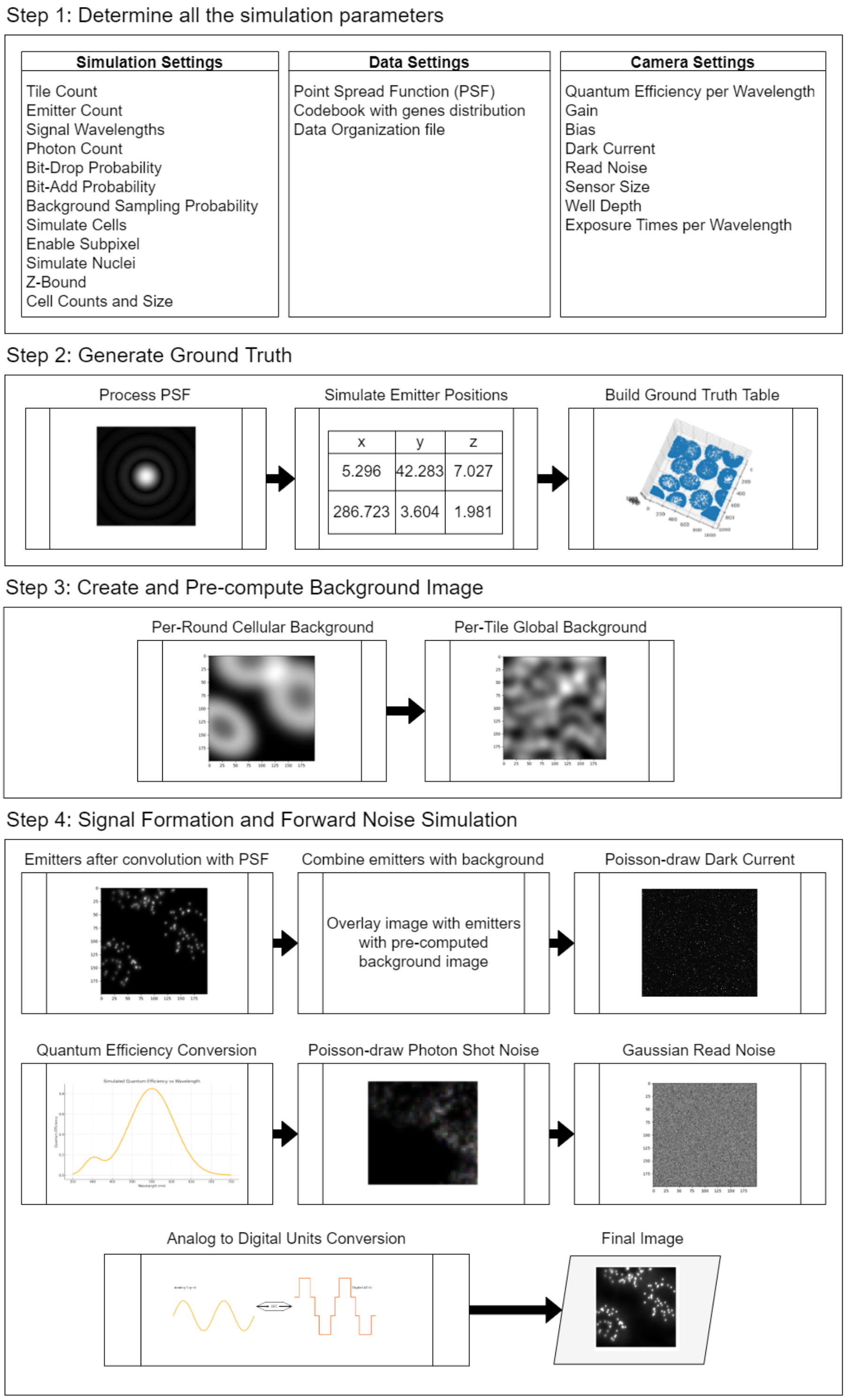


**Supplemental Figure 9**: **MERFISH image simulation pipeline.** The workflow comprises four sequential stages. **(1)** Parameterization: user-defined simulation settings (tile count, emitter count, photon count, bit-add/drop and background sampling probabilities), data inputs (PSF, codebook, data organization) and camera characteristics (quantum efficiency, gain, bias, dark current, read noise, sensor geometry, exposure times). **(2)** Ground-truth generation: processing of the PSF, stochastic placement of emitters in 3D and assembly of a comprehensive ground-truth table. **(3)** Background modeling: synthesis and pre-computation of per-round cellular backgrounds and per-tile global backgrounds. **(4)** Signal formation and noise simulation: PSF convolution of emitters, overlay onto background, Poisson sampling of dark current and photon shot noise, quantum efficiency conversion, addition of Gaussian read noise and analog-to-digital conversion.

**Supplementary Table 5**: **Noise and specificity metrics using off-target biological controls.** Evaluation of false-positive transcript assignments by measuring the detection of T/B cell marker genes, which are expected to be silent in the NRG host tissue. CoSiD substantially reduces the off-target expression fraction and yields higher signal-to-noise ratios (SNR) relative to MERlin.

| | **CoSiD** | **MERlin** |
|---|---|---|
| **Off-Target Signal (T/B Markers)** | | |
| Off-target fraction (TB genes) | 0.0009 | 0.0064 |
| Cells expressing TB genes (%) | 18.4493 | 57.4717 |
| **Signal-to-Noise Ratio** | | |
| SNR (targets vs blanks) | 1.3309 | 1.0846 |
| SNR (targets vs TB genes) | 1.2351 | 0.7709 |
| **Mutual Exclusivity Co-expression** | | |
| MECR: Neutrophil vs Macrophage | 0.0648 | 0.1328 |
| MECR: Neutrophil vs Endothelial | 0.0490 | 0.0958 |
| MECR: Endothelial vs Macrophage | 0.2420 | 0.2406 |
| MECR: Mean | 0.1186 | 0.1564 |

TB genes (*Cd28, Stat4, Il21r, Icos, Ms4a1, Cr2, Cd19*) are panel-included probes expected silent in NRG host tissue. SNR = $\log_{10}$(mean target / mean control). MECR = co-expression rate of mutually exclusive marker pairs.

**Supplementary Table 6**: **Global False Discovery Rate (FDR) estimation.** FDR computed using panel-included non-targeting blank barcodes as proxies for decoding noise.

| | **CoSiD** | **MERlin** |
|---|---|---|
| **Detection Counts** | | |
| Blank detections | 1,020 | 4,801 |
| Gene molecule detections | 5,878,085 | 4,139,278 |
| Total molecule detections | 5,879,105 | 4,144,079 |
| N blank features | 6 | 6 |
| N gene targets | 134 | 134 |
| **False Discovery Rate** | | |
| Global FDR | $3.87 \times 10^{-5}$ | $2.59 \times 10^{-4}$ |

Global FDR = (N_blank / N_total) × (G_genes / G_blank) × 1/100. Blank features are panel-included non-targeting control barcodes. Formula follows Plummer et al., *Nat. Biotechnol.* 2025.

**Supplementary Table 7**: **Noise and specificity metrics for the MERFISH 4T1 cell culture dataset.** Evaluation of off-target transcript assignments using T/B cell marker genes, which are expected to be biologically silent in the pure 4T1 breast cancer cell culture. Performance is compared across CoSiD, the default MERlin decoder, and a matched scRNA-seq reference.

| | **CoSiD** | **MERlin** | **scRNA-seq** |
|---|---|---|---|
| **Off-Target Signal (T/B Markers)** | | | |
| Off-target fraction (TB genes) | 0.0019 | 0.0028 | 0.0001 |
| Cells expressing TB genes (%) | 65.4691 | 74.0260 | 2.5966 |
| **Signal-to-Noise Ratio** | | | |
| SNR (targets vs blanks) | 1.7643 | 1.6610 | – |
| SNR (targets vs TB genes) | 1.3466 | 1.2142 | 1.6796 |

TB genes (*Cd28, Stat4, Il21r, Icos, Ms4a1, Cr2, Cd19*) are panel-included probes expected silent in 4T1 cells. SNR = $\log_{10}$(mean target / mean control). MECR omitted: no distinct cell types expected in cell line data.

**Supplementary Table 8**: **Standardized spatial quality control metrics for the DART-FISH brain dataset.** Performance metrics evaluating transcript recovery and signal specificity across three decoding algorithms: original sparse deconvolution (SpD), CoSiD, and MERlin. Metrics were computed using the SpatialQM framework on nuclear-restricted transcripts to minimize co-detection artifacts.MECR was computed using excitatory vs inhibitory neuronal markers.

| **Standardized Spatial QC Metrics** | | | | | | | |
|---|---|---|---|---|---|---|---|
| **Decoder** | **N Cells** | **Transcripts/Nuc** | **Transcripts/Area** | **Sparsity** | **Entropy** | **MECR** | **Global FDR** |
| SpD | 25,597 | 25.453 | 0.019 | 0.921 | 0.571 | 0.216 | – |
| CoSiD | 26,279 | 41.416 | 0.032 | 0.836 | 0.960 | 0.327 | 0.0027 |
| MERlin | 26,273 | 41.483 | 0.032 | 0.833 | 0.964 | 0.325 | 0.0031 |

Metrics computed using SpatialQM framework [35]
MECR = Mutual Exclusivity Co-expression Rate; FDR = Global False Discovery Rate.

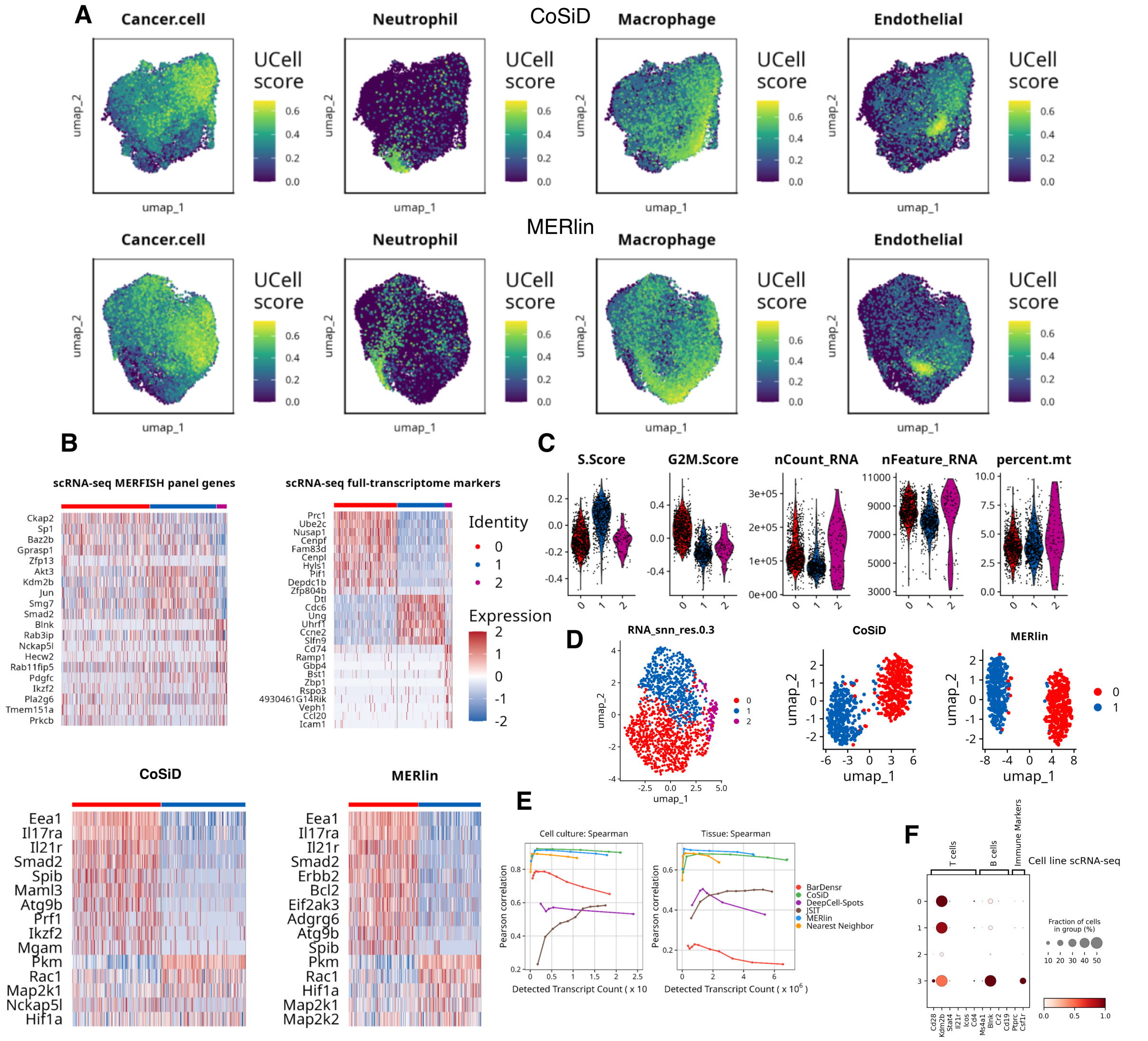

**Supplemental Figure 10**: **Characterization of the 4T1 cell culture and tissue heterogeneity** **(A)** UCell Signature scores from the scGate classification applied to the 4T1 tissue where cells with a positive score > 0.2 and no competing singaure exceeding 0.2 were assigned to that type. **(B)** Heatmap visualizing differential expression testing of the scRNA-seq 4T1 cell culture using either the full transcriptome or targeted MERFISH panel genes. **(C)** Per-cell distributions of cell cycle scores and QC metrics for scRNA-seq 4T1 cell culture. **(D)** UMAPs comparing clustering of scRNA-seq (left) and decoded outputs (right) using the target panel genes. **(E)** Spearman correlations as a function of detected transcript counts in the 4T1 cell culture dataset (left) and 4T1 tissue (right), comparing six decoding methods (MERlin, Nearest Neighbor, CoSiD, BarDensr, DeepCell-Spots, and JSIT). **(F)** Dot plot of the pre-filtered scRNA-seq cell culture, visualizing off-target immune marker genes.

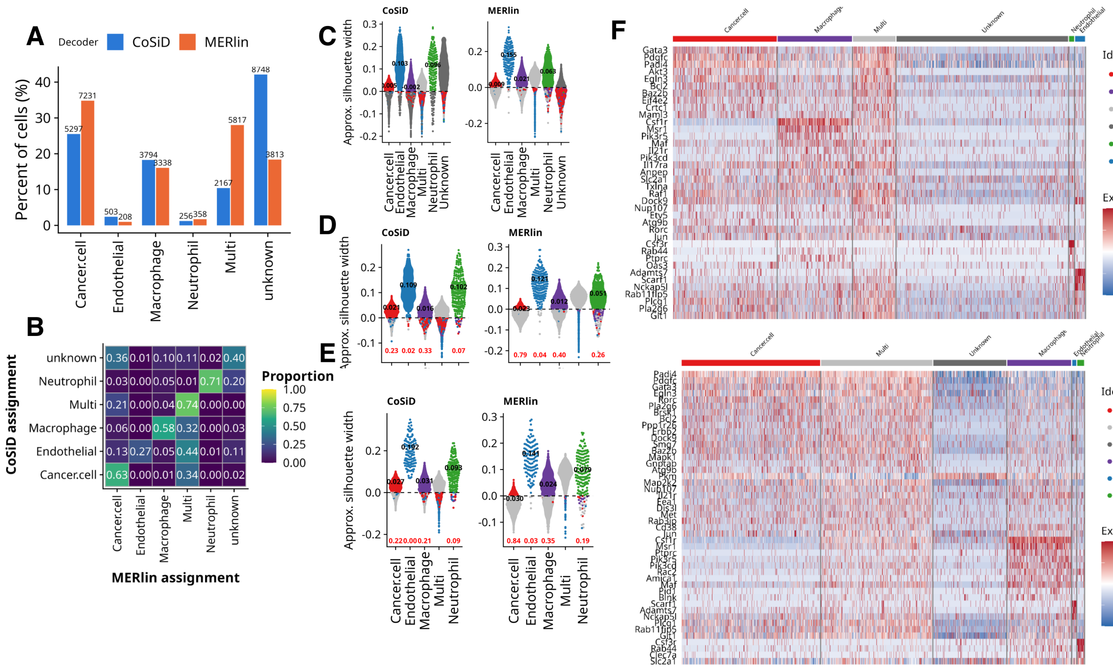


**Supplemental Figure 11**: **Comparison of cell-type annotation and cluster separation between CoSiD and MERlin decoding in the 4T1 tissue.** **(A)** Proportion of cells assigned to each scGate class in the 4T1 tissue dataset under CoSiD and MERlin decoding. **(B)** Heatmap showing concordance of per-cell assignments between the decoders. Each cell is cross tabulated by its CoSiD label(rows) and MERlin label (columns), values row-normalized to the CoSiD total. **(C-E)** Approximate silhouette width (bluster::approxSilhouette, PCA embedding) for CoSiD (left) and MERlin (right), with per-group median overlaid in black text and negative cell fractions indicated in red text. The three panels differ only in which cells participate in the computation. **(C)** All groups included, with unknown and Multi treated as clusters. **(D)** Unknown cells removed and PCA recomputed on the remaining cells. **(E)** PCA computation restricted to the [7,476] cells assigned the same cell type by both decoders (diagonal in B). **(F)** Expression heatmaps of panel genes across annotated groups for CoSiD (top) and MERlin (bottom).

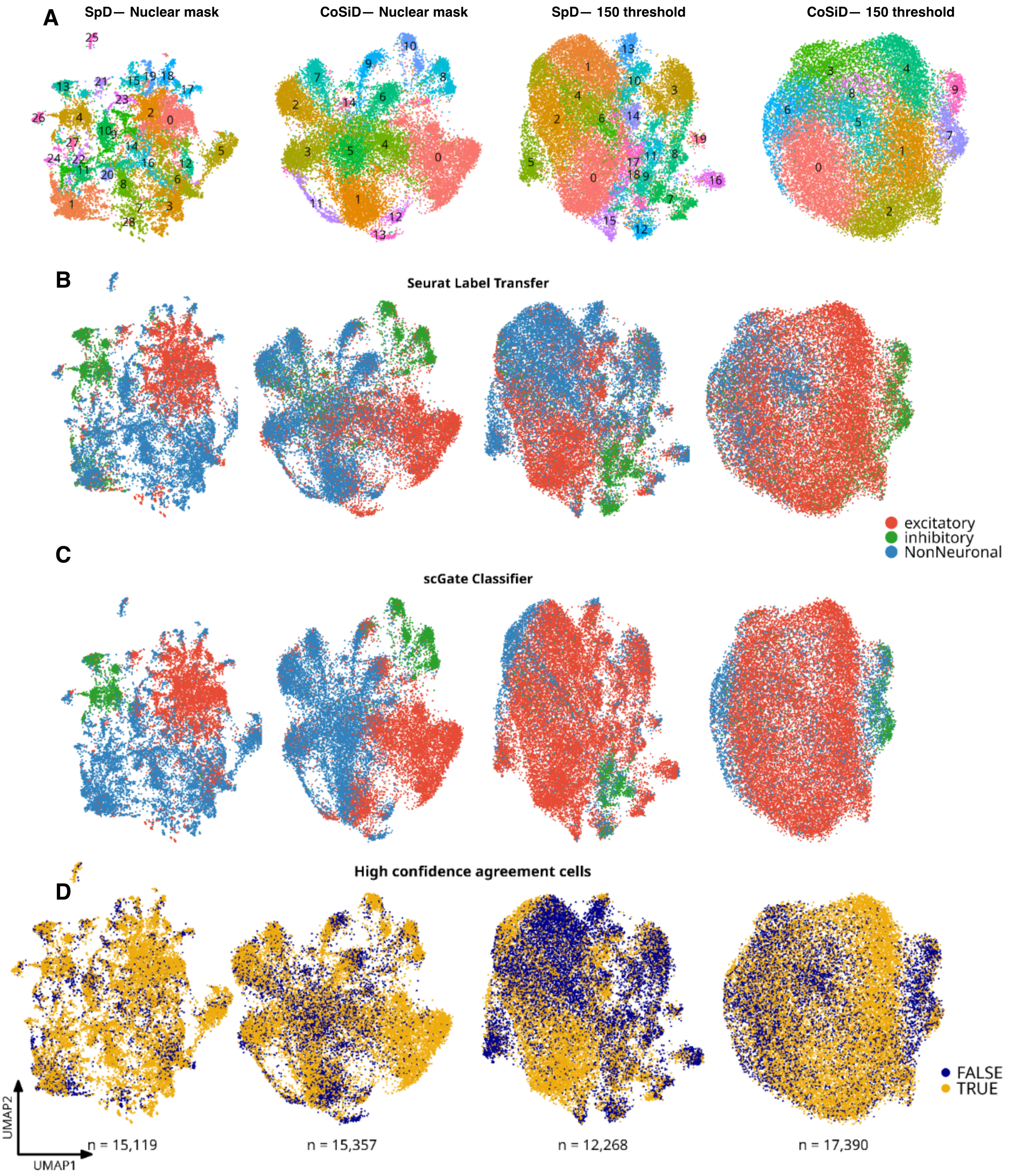


**Supplemental Figure 12**: **Impact of transcript assignment thresholds on DART-FISH clustering and annotation.** **(A)** UMAP visualizations of unsupervised clusters across decoded datasets and thresholding conditions, illustrating differences in clustering structure between SpD and CoSiD using stringent nuclear masks or a maximum threshold of 150 set by the SpD software parameter. **(B)** Major-class annotations obtained by Seurat label transfer from the matched snRNA-seq reference. **(C)** Independent major-class annotation using the scGate marker-based classifier. **(D)** High-confidence cells defined by concordant major-class assignments between Seurat label transfer and scGate.

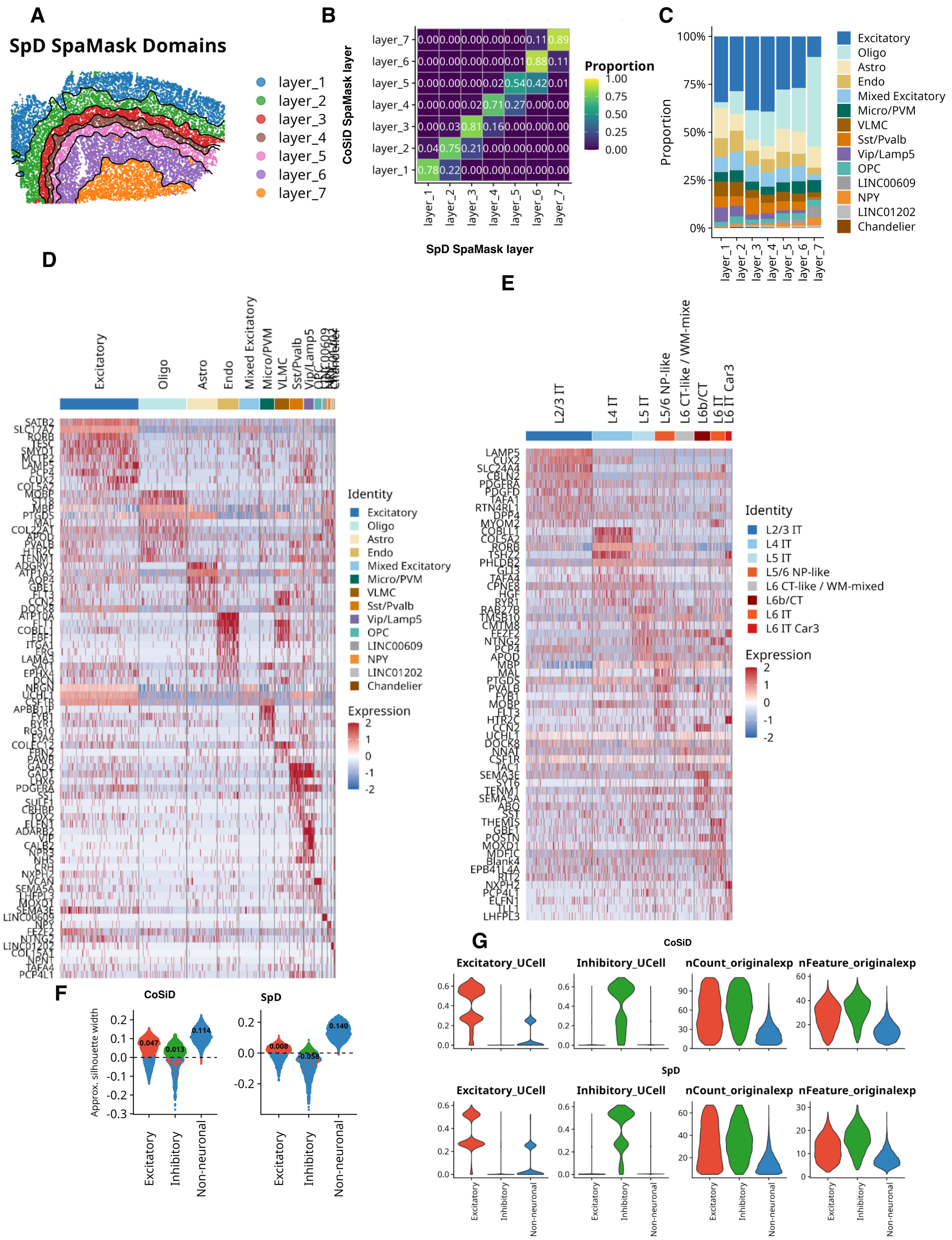


**Supplemental Figure 13**: **Spatial domain and cell-type characterization of the DART-FISH brain dataset.** **(A)** SpaMask-inferred spatial domains from the SpD-decoded DART-FISH dataset. **(B)** Heatmap showing overlap between SpaMask layers inferred from CoSiD-decoded and SpD-decoded data. **(C)** Composition of CoSiD-derived SpaMask layers across all annotated cell types, illustrating spatial enrichment of neuronal and non-neuronal populations across cortical layers. **(D)** Differential expression heatmap of marker genes across all annotated CoSiD-derived cell classes, detailing major-class and subclass-level annotation of excitatory, inhibitory, and non-neuronal populations. **(E)** Differential expression heatmap of marker genes across CoSiD-derived excitatory neuron subclasses, supporting annotation of superficial, intermediate, and deep excitatory populations. **(F)** Approximate silhouette width by cell class for CoSiD and SpD decoders. Each point represents one cell; median silhouette width per class is overlaid. **(G)** Violin plots showing the distribution of excitatory and inhibitory UCell signature scores, transcript counts, and detected genes per cell for CoSiD and SpD.
